# Virome analysis of irrigation water sources provides extensive insights into the diversity and distribution of plant viruses in agroecosystems

**DOI:** 10.1101/2023.06.20.545696

**Authors:** Olivera Maksimović, Katarina Bačnik, Mark Paul Selda Rivarez, Ana Vučurović, Nataša Mehle, Maja Ravnikar, Ion Gutiérrez-Aguirre, Denis Kutnjak

## Abstract

Plant viruses pose a significant threat to agriculture. Several are stable outside their hosts, can enter water bodies and remain infective for prolonged periods of time. Even though the quality of irrigation water is of increasing importance in the context of plant health, the presence of plant viruses in irrigation waters is understudied. In this study, we conducted a large-scale high-throughput sequencing (HTS)-based virome analysis of irrigation and groundwater sources to obtain complete information about the abundance and diversity of plant viruses in such waters. We detected nucleic acids of plant viruses from 20 families, discovered several novel plant viruses from economically important taxa, like *Tobamovirus* and observed the influence of the water source on the present virome. By comparing viromes of water and surrounding plants, we observed presence of plant viruses in both compartments, especially in cases of large-scale outbreaks, such as that of tomato mosaic virus. Moreover, we demonstrated that water virome data can extensively inform us about the distribution and diversity of plant viruses for which only limited information is available from plants. Overall, the results of the study provided extensive insights into the virome of irrigation waters from the perspective of plant health. It also suggested that an HTS-based water virome surveillance system could be used to detect potential plant disease outbreaks and to survey the distribution and diversity of plant viruses in the ecosystems.

## 1. Introduction

Plant viruses are a well-known risk factor in crop production, resulting in at least $30 billion in yield losses annually [1]. The presence of plant viruses in the aqueous environment has been known for nearly four decades [2]. They have been detected in a variety of water bodies including rivers, lakes, ice, and tap water [3], and in wastewaters [4]–[7]. For some of them, the stability in water for prolonged period and the ability to infect plants after this time were demonstrated [8]. Although the research of plant viruses in environmental waters has been gradually developing, there are still relatively few studies focusing on this topic. There are even fewer studies that focused on irrigation water and/or surface water near farms [9]–[11]. In one study, researchers from China have confirmed presence and infectivity of eight selected tobamoviruses [12]. These findings put forward possible risks associated with the use of water containing plant viruses for irrigation of plants, which need to be further investigated. This is especially important since currently, 70% of the freshwater withdrawals are used for irrigation and general agricultural needs and this trend is expected to increase [13]. For example, in the European Union, the main irrigation water resource is on-farm underground water or surface supply networks in more arid areas like Greece [13]. In addition to these two sources, on-farm surface water (e.g., rivers, streams, ponds) is also often used [13].

In recent decade, virome studies, based on high-throughput sequencing (HTS), enabled high resolution generic studies of virus diversities in environmental water samples. Many studies addressed diversity of viruses (e.g., bacteriophages) in oceans [14], [15], however, less attention has been given to freshwater bodies [16]. Virome studies of freshwater bodies, specifically focusing on plant viruses are sparse [9], [17], even though this aspect is important in the view of the possible detrimental effects of plant viruses in agriculture, if such waters are used for irrigation of plants [3], [8]. To better understand the potential risks associated with the presence of plant viruses in irrigation waters, baseline virome studies are first needed to provide information about the presence of viruses in such samples.

Moreover, large sequence datasets obtained by HTS-based analysis of environmental water samples can bring additional information, reaching beyond the presence/absence data for different viruses. This was most evidently demonstrated during the COVID-19 pandemic where wastewater monitoring has been used for tracking virus variants in populations across countries [18]. Extending such framework to other water types could bring extensive information about different aspect of epidemiology of other viruses. For example, environmental monitoring of water used in, and around agricultural sites could be used to detect and anticipate the entry and spread of plant viruses in the given area and to better understand their epidemiology.

In this study, we investigated the virome of diverse types of irrigation water and nearby environmental ground water samples, collected in several agroecosystems (tomato farms) in Slovenia. Focusing on plant viruses, we aimed to (1) provide a baseline virome data for plant virus presence in sampled waters, (2) compare viromes of different irrigation water types,(3) use water virome data to discover novel plant viruses, and (4) obtain information about plant virus diversity and distribution in the wider environment. Moreover, these results were associated and compared with results from a previous study [19] that looked at the virome of tomato and surrounding weed plants at the same tomato farms. This enabled us to obtain unique and unprecedented comparative insight into the virome compositions of two different but connected agroecosystem components: plants and water.

By elucidating the presence, diversity, and distribution of plant viruses in irrigation water, we can better understand their potential impacts on agricultural systems and develop strategies for their early detection and management. The applicability of water analysis for detection of new and/or emergent viruses is also discussed herewith, following its usability as an early warning system for viruses just entering the environment. This research contributes to the growing body of knowledge on plant viruses in environmental waters and highlights the need for further studies in this area that would improve our understanding of the ecological dynamics of plant viruses in agroecosystems.

## 2. Materials and Methods

### 2.1. Samples and sampling locations

Water samples (5 L) were collected in the summer of 2019 and 2020 at different locations in Slovenia (Figure 1, Supplementary Information 1, S1). Selected locations were farms with tomato as the main growing crop. The sampled water was either used at the farm to irrigate crops or from the nearby groundwater body not primarily used for irrigation. Water was collected in autoclaved glass bottles and transported back to the laboratory in cooler boxes. Before further processing, samples were stored at 4 ℃ for up to 48 hours. Different types of samples are labelled further in the text as: (T) - tap water from municipal water system; (U) - underground water originating from any underground source; (P) - pond or any standing freshwater body, and (R) - rivers or any type of ground watercourse, including canals and streams. The sample nomenclature shows the type of water, year of sampling, and site number (e.g., U-19-01 represents underground water sampled in 2019, at location 1).

**Figure 1.**
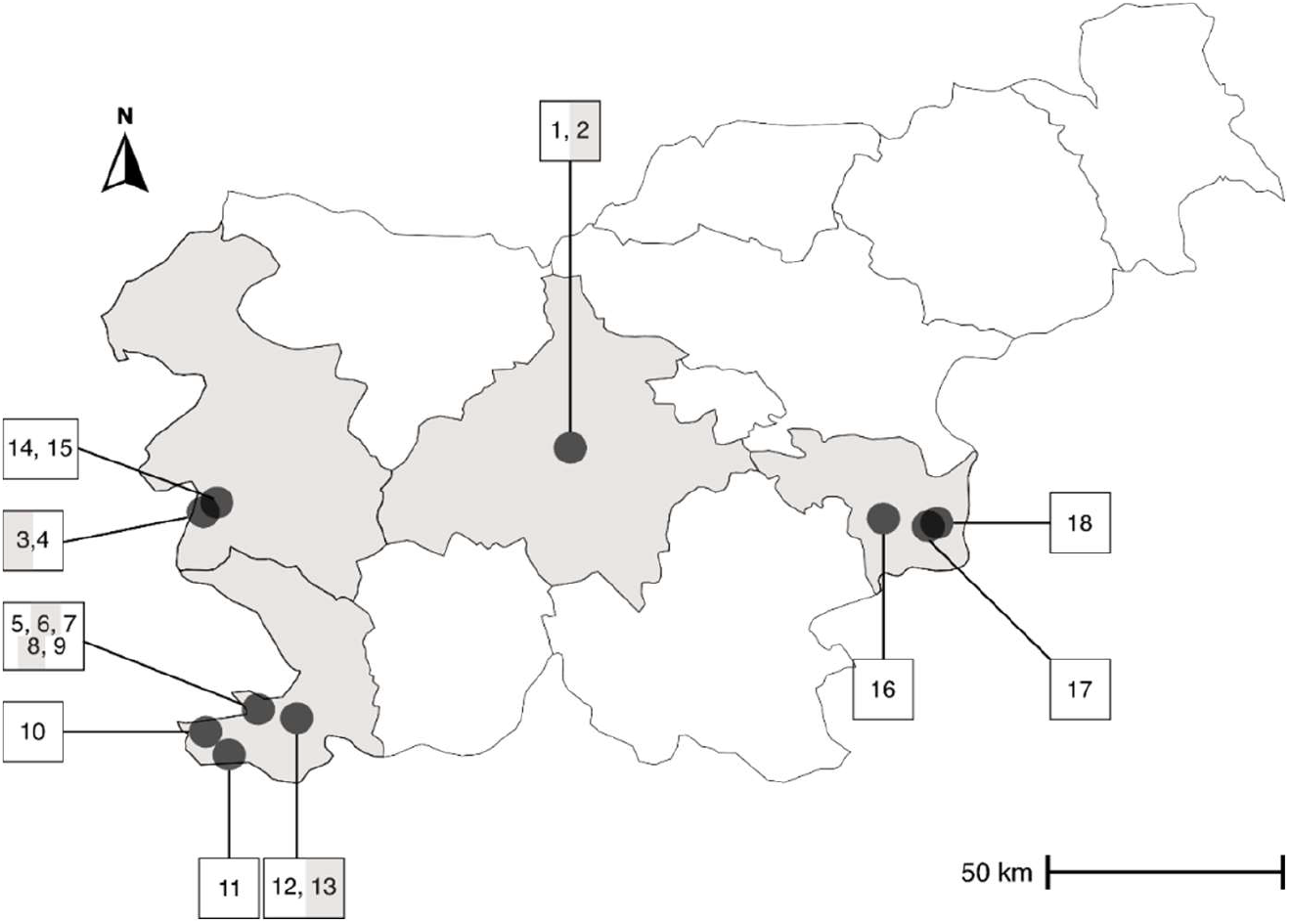
Sampling locations and their geographical distribution. Dots represent sampling location (if locations were very close together, they are designated with the same dot). Lines separate different statistical regions. Greyed-out regions contain sampling locations. Grey shaded numbers represent locations where ground water not directly used for irrigation was sampled. For exact longitude and latitude information refer to Supplementary Information 1, S1.

### 2.2. Concentration of water samples and nucleic acids extraction

All samples (5L) were concentrated using convective interaction media (CIM) monolithic chromatography, with a CIM quaternary amine (QA) 8 mL column (Sartorius BIA separations, Slovenia), and with step gradient elution to a final elution volume of 25 mL. The collected fractions (one before concentration (raw) and one after concentration (elution) for each sample) were stored at −80 °C until nucleic acids extraction. RNA from elution fraction of each sample was extracted using two different protocols (QIAmp Viral RNA Mini kit (Qiagen, USA) and TRIzol LS (Invitrogen, USA)), and RNA from raw fraction only with QIAmp Viral RNA kit, with each batch accompanied by negative control of isolation (NCI, nuclease-free water used instead of sample). The detailed protocol is available in a previous published study [4]. Before extraction using the QIAmp Viral RNA Mini Kit, samples, including NCI were spiked with 2 ng of Luciferase Control RNA (LUC) (Promega, USA). These extracts were used for real-time quantitative PCR (RT-qPCR) analysis. TRIzol LS extracts were used for sequencing, wherein only the NCI was spiked with LUC RNA. The RNA extracts were stored at −80°C until further analysis.

### 2.3. RT-qPCR

RNA extracted using QIAamp Viral RNA kit from raw and elution fraction were analysed using RT-qPCR for the targeted detection of three selected tobamoviruses, namely, pepper mild mottle virus (*Tobamovirus, Virgaviridae,* PMMoV), tomato mosaic virus (*Tobamovirus, Virgaviridae,* ToMV), and cucumber green mild mottle virus (*Tobamovirus, Virgaviridae,* CGMMV). For PMMoV and ToMV, both raw and elution fraction were tested, and for CGMMV, only the elution fraction was tested by RT-qPCR. In addition, an RT-qPCR assay specific for luciferase RNA was used to quantify LUC RNA that was spiked during the nucleic acid extraction step. Published primer/probe sets were used for all targets (PMMoV [20], ToMV [11], CGMMV [21] and LUC [22]). Each RT-qPCR reaction was performed with 2 µL of extracted RNA per reaction in a total reaction volume of 10 µL. RT-qPCR was performed using the AgPath-ID™ One-Step RT-qPCR Kit (Life Technologies, USA) on a 7900HT Fast Real-Time PCR System (Applied Biosystems, USA), using cycling parameters as recommended by the mastermix manufacturer. Samples were tested in duplicates and prepared as undiluted and 10-fold dilutions. A negative template control (nuclease-free water instead of the RNA) and a positive control (with known presence of the corresponding target virus) were included for each assay in each RT-qPCR analysis. Data were analysed in standalone software (SDS v4.0) with automatic setting of the baseline and threshold set up to a value of intersection between amplification curves at the exponential phase of the amplification, namely, 0.065 for PMMoV, 0.02 for ToMV, 0.15 for CGMMV and 0.15 for LUC assays. Each amplification plot was checked manually, and the result was considered positive if it produced an exponential amplification curve distinguishable from negative controls. In such cases, Cq values were calculated. Cq values for LUC RNA were monitored in a control chart and extraction was considered successful if obtained Cq was within ±3 standard deviations from the mean Cq value (data not showed).

### 2.4. Shotgun high-throughput sequencing (HTS)

RNA isolates of each sample, along with 3 spiked NCIs (spiked with LUC RNA), all obtained by the TRIzol LS extraction protocol, were randomly pre-amplified according to the protocols described previously [4]. The pre-amplification products were sent to SeqMatic LLC (Fremont, USA) for library preparation and sequencing. Nextera XT DNA Library Prep Kit (Illumina, USA) was used to prepare the sequencing libraries, which were shotgun-sequenced using an Illumina MiSeq platform (Illumina, USA) in a 2×250 bp mode.

### 2.5. Analysis of HTS data

#### 2.5.1. Data preparation

After obtaining the raw sequencing data, sequencing reads were trimmed-off of sequencing adaptors and primer sequences from the pre-amplification step and filtered with a quality filter (Supplementary Information 1, S3) in CLC Genomics Workbench (CLC-GWB) (v. 20-22). Datasets were then normalized by random subsampling of each sample with the subsampling size equivalent to the lowest number of reads observed among the samples within the same year (Supplementary Information 1, S1).

#### 2.5.2. Virome analysis focusing on plant viruses

A general overview of metagenome and detection of known viruses was done by first exporting normalized reads subsets from CLC-GWB (v. 20-22) and comparing them for similarity to the entire NCBI nr database (v. 237) using DIAMOND (v. 9.34) blastx [23] with default parameters. The results of the DIAMOND similarity searches were used as input for the taxonomic classification of reads using MEGAN (Metagenome Analyzer, v. 6.20.16, May 2020 database) [24] with the LCA algorithm (Supplementary Information 1, S3). The obtained MEGAN outputs in the form of summarized reads were used to provide an overview of the taxonomic classification of the sequencing reads (Supplementary Information 1, S4). Additionally, viral reads binned on the level of order were used to determine the genome organisation (as denoted by International Committee on Taxonomy of Viruses (ICTV) for each order [25]) and viral reads binned on the level of family were used to determine the expected host (as denoted by ICTV for each family [25]), data in Supplementary Information 1, S5, S6. Read classifications on different levels were visualized as bar plots using RStudio (v. 2021.09.0) and edited in Inkscape (v. 0.92).

Information on classification of reads for each plant virus genus was exported from MEGAN as a BLAST table (Export→Matches). The table was further curated using a custom script (Supplementary Information 2) which classified reads that were assigned to a specific species up to the level of genus if they were below the ICTV [25] species demarcation criteria for that genus. Data generated in this manner for each genus is summarized in Supplementary Information 1, S7. This way, we obtained the number of reads assigned per virus species and genera for each sample. Reads classifications for plant virus families, genera and species were visualized as bubble charts in RStudio (v. 2021.09.0) and edited in Inkscape (v. 0.92).

#### 2.5.3. Assembly of genomic sequences for new plant viruses

Performed read classifications showed the possible presence of new plant virus species (e.g., reads classified only at the genus level). Thus, we aimed to assemble genomic sequences of new viruses using two different approaches. In the first approach, all reads, assigned to the genus level for plant virus genera, which had over 100 reads assigned in MEGAN, were *de novo* assembled in CLC-GWB (v. 20-22) (parameters in Supplementary Information 1, S3). After assembly, any contig longer than 500 bp was checked using BLASTx against an entire NCBI nr database (v. 248). Contigs that did not match to a known virus and with percent identity above the species demarcation criterium (as listed by the ICTV [25]) for that genus were kept for further analysis. Such contigs were extended in Geneious Prime (v. 2022.2) by iterative mapping of reads from the corresponding sample back to the contig until there was no more extension (parameters in Supplementary Information 1, S3). Finally, all relevant contigs were checked for open reading frames (ORFs) in CLC-GWB (v. 20-22) to confirm the genome structure of complete or near-complete genomes of novel viruses.

In the second approach, all reads (after normalisation) per sample were *de novo* assembled using SPAdes (v. 3.14) [26], and compared for similarity to an entire NCBI nr database (v. 237) using DIAMOND (v. 9.34) blastx [23] with default parameters. The results of the DIAMOND alignment were used as an input for the taxonomic classification of reads using MEGAN (v. 6.20.16, May 2020 database) [24]. Contigs were manually inspected, and those matching plant viruses on the level of genus, but not matching a known virus on the level of species (based on the percent identity species demarcation) were extended in Geneious Prime (v. 2022.2) and checked for ORFs as described above.

#### 2.5.4. Pairwise identity comparisons and phylogenetic analyses for putative new plant virus species

For each genus that contained a putative novel virus, complete genome sequences from selected members of the genus from the NCBI GenBank database and the sequences of the potential new viruses from this study were aligned in MEGA X (v. 10.0.5) [27] using ClustalW with default parameters. These alignments were used in SDT (v. 1.2) [28] to calculate pairwise identities using the software’s MUSCLE algorithm and were visualized in RStudio. Sequences of potentially novel viruses for which pairwise identities were below the species demarcation criteria for a corresponding genus (ICTV [25]) were further analysed.

Phylogenetic trees were constructed using the amino acid sequence of RNA-dependent RNA polymerase (RdRp) gene for selected known member species of that genus, new virus species discovered in this study, and an outgroup virus. The only exception is for *Sobemovirus* genus, where the complete genome nucleotide sequences were used instead. Outgroup viruses were selected from a different genus of the same virus family, following ICTV [25] recommendations, where possible. The selected sequences were aligned in CLC-GWB (v. 20-22, Supplementary Information 1, S3), and the most conserved region was selected for phylogenetic analysis (Supplementary Information 1, S8). This alignment was further trimmed using the ‘automated1’ method in trimAl (v. 1.3) [29]. Phylogenetic trees were constructed using IQtree (v. 1.6.12) using the maximum-likelihood approach, with the selection of the most appropriate substitution models performed in the same software [30] (Supplementary Information 1, S9). The phylogenetic trees were visualised in iTOL (v. 6.7) [31].

#### 2.5.5. Comparison of water viromes and viromes of surrounding plants

In a parallel study [19] samples from tomatoes and surrounding weed plants, from the same locations as in this study, were analysed for the presence of plant viruses. In order to compare viruses that were found in these plants to the viruses found in water samples, reads from each water sample were mapped using CLC-GWB (v. 20-22, parameters in Supplementary Information 1, S3) to a user-defined database consisting of all viral sequences detected in the analysed plants (a complete list of viruses detected in plants can be found in [19]). Occurrence of viruses detected both in water and plants, and abundance of reads corresponding to selected viruses found in water, were plotted on the map of the sampling locations using RStudio (v. 2021.09.0) and edited in Inkscape (v. 0.92).

#### 2.5.6. Analysis of the genomic diversity of Plantago tobamovirus 1 in water and plant samples

Plantago tobamovirus 1 (*Tobamovirus, Virgaviridae*) (PTV1) was detected in *Plantago major* for the first time in a plant [19] at one of the sampled locations. Subsequent analysis showed its presence in water samples in various other locations covered in the study. To explore the variability of PTV1, contigs (generated using the second assembly approach, Section 2.5.3) from all samples that had > 90% identity with PTV1 in BLASTn analysis against an entire NCBI nr database (v. 248) were aligned in CLC-GWB (V. 20-22, parameters in Supplementary Information 1, S3). Two longer genome parts in which several contigs overlapped, were used for further analysis (Supplementary Information 1, S9). Pairwise identities based on those two alignments were calculated using SDT (v. 1.2) [28]. In the next step, the two selected alignments were additionally aligned using the same algorithm to include ribgrass mosaic virus (*Tobamovirus, Virgaviridae*) as an outgroup. Phylogenetic trees were constructed from those assemblies using neighbour joining method in CLC-GWB (v.20-22, parameters in Supplementary Information S3, S9) and the leaves representing different consensus virus genomes were connected to corresponding locations on the map, using Inkscape (v. 0.92).

## 3. Results

### 3.1. Targeted detection confirms the efficiency of concentration approach and shows the presence of nucleic acids of pathogenic plant viruses in analysed water samples

Each water sample was concentrated using the method described to increase the relative abundance of present viruses. The efficiency of concentration was tested by targeted qPCR for two selected viruses expected to be present in many samples. The method performed efficiently as seen from the reduction of Cq values from raw to elution fraction for PMMoV and ToMV (Supplementary Information 3, Figure 9). In the cases where Cq reduction could be calculated, it ranged from 1.8 to 8.6. In addition, CGMMV was tested only in the elution fraction (Supplementary Information 1, S2). The Cq values obtained for the detected viruses (elution fraction) varied between Cq 36 and 20. ToMV was most abundantly present and detected in 21 out of 22 samples, followed by CGMMV (18/22) and PMMoV (15/21).

### 3.2. The metagenome overview and the effect of the water source on the amount and diversity of detected plant virus sequences

Taxonomic classification of reads was performed, to obtain a general overview of the metagenome of the concentrated water. A considerable proportion of reads (37.6-80.5%, depending on the sample) did not provide matches in the used database. Bacteria accounted for the second largest proportion (5.9-58.8%), followed by eukaryotes (1.3-44.5%). As for viral reads, they account for 0.01 to 7.7% reads per sample (Figure 2a). Across the samples, we detected reads corresponding to all the genome organization types for viruses, with (-) ssRNA and dsRNA viruses being the least abundant (< 1% of all viral reads for samples in which they were detected). We further focused on ssRNA viruses, since 75% of known plant viruses have this genome organization [32]. Per sample, between 0.6% and 95.6% of reads assigned to viruses were indeed classified as ssRNA viruses. In addition, unclassified RNA viruses (which can have any variation of RNA genome organization) were the second most abundant group, accounting for 0.3-60.4% of viral reads per sample (Figure 2b). High abundance of plant virus sequences was confirmed by looking at the number of viral reads binned by the expected host organism. In this analysis, in almost all samples, majority of viral reads were assigned to viruses infecting bacteria and archaea (when excluding the wide host range/unclear category), followed by viruses infecting plants, fungi and protists. In the case of samples P-19-03, U-19-01 and R-19-13, plant/fungi/protist viruses are the most abundant subset (Figure 2c).

**Figure 2.**
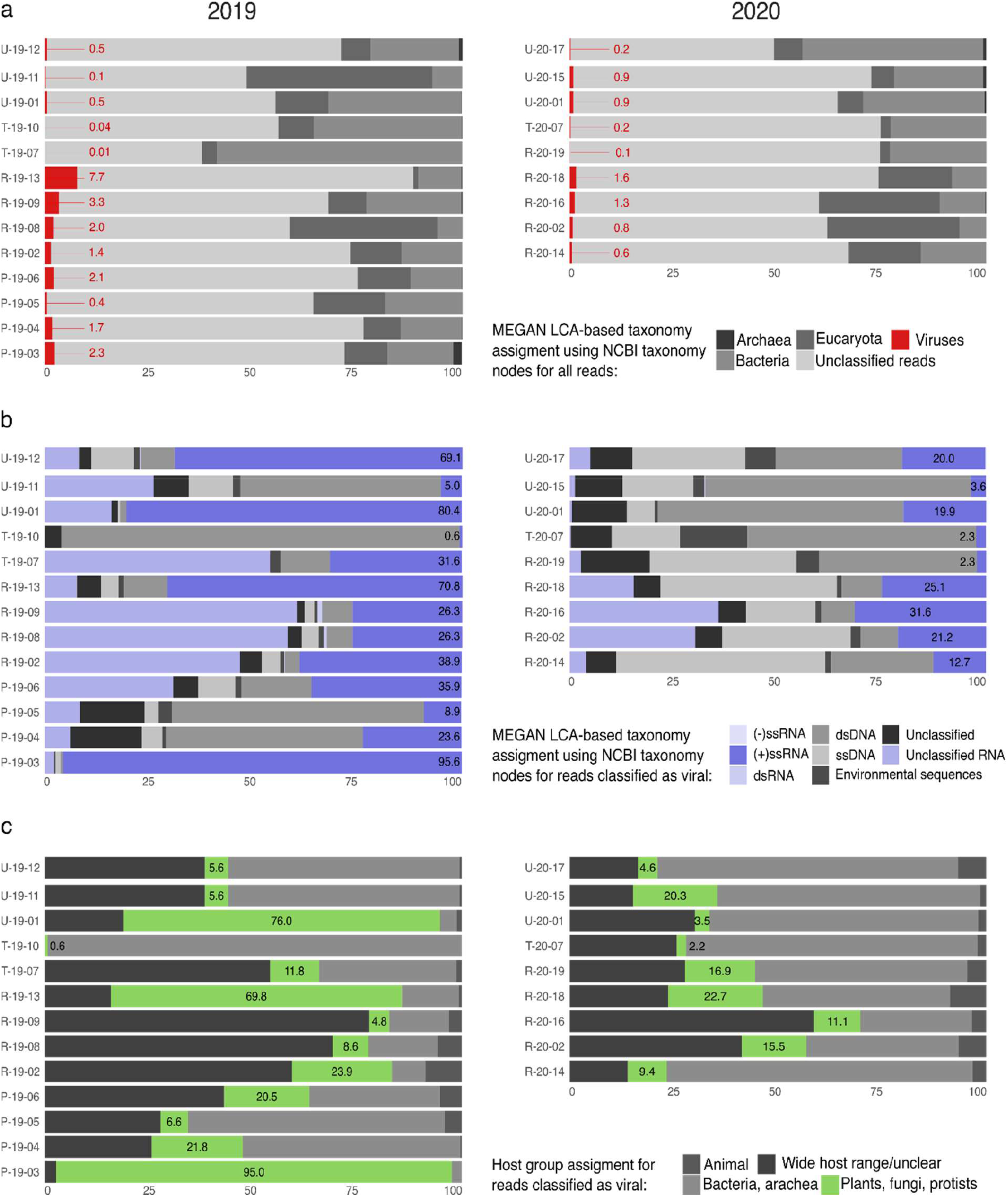
Classification of sequencing reads for different collected water samples, based on DIAMOND blastx similarity search, followed by MEGAN-LCA binning. (a) “Domain” level classification of all reads – highlighted and noted in red are proportions (%) of reads belonging to viruses. (b) Classification of viral reads according to their genome types based on the “order” level classification associated with available genome type information from ICTV – highlighted in shades of purple are proportions (%) of viral reads belonging to RNA viruses with different genome types. Numbers denote relative proportion (%) of (+)ssRNA viral reads. (c) Classification of viral reads according to predicted host groups based on the “family” level classification associated with available host information from ICTV [25] database – denoted and highlighted in green are proportions (%) of reads belonging to host group “Plants, fungi and protists”. In all panels, samples are separated in two columns by year.

Plant viruses from a total of 20 families were detected across samples (Figure 3). The most abundant viruses belonged to the *Virgaviridae* and *Tombusviridae* families, for which reads were detected in 19 out of 22 analysed samples (Figure 3). We can observe a difference between groundwater and underground water samples subgroups at this taxonomic level. For the groundwater samples, the number of families detected per sample ranged from 2 to 17, with the average number being 10. On the other hand, for the underground samples, the average number of detected families is 5 and in none of the samples we detected more than 9 families (Figure 3).

**Figure 3.**
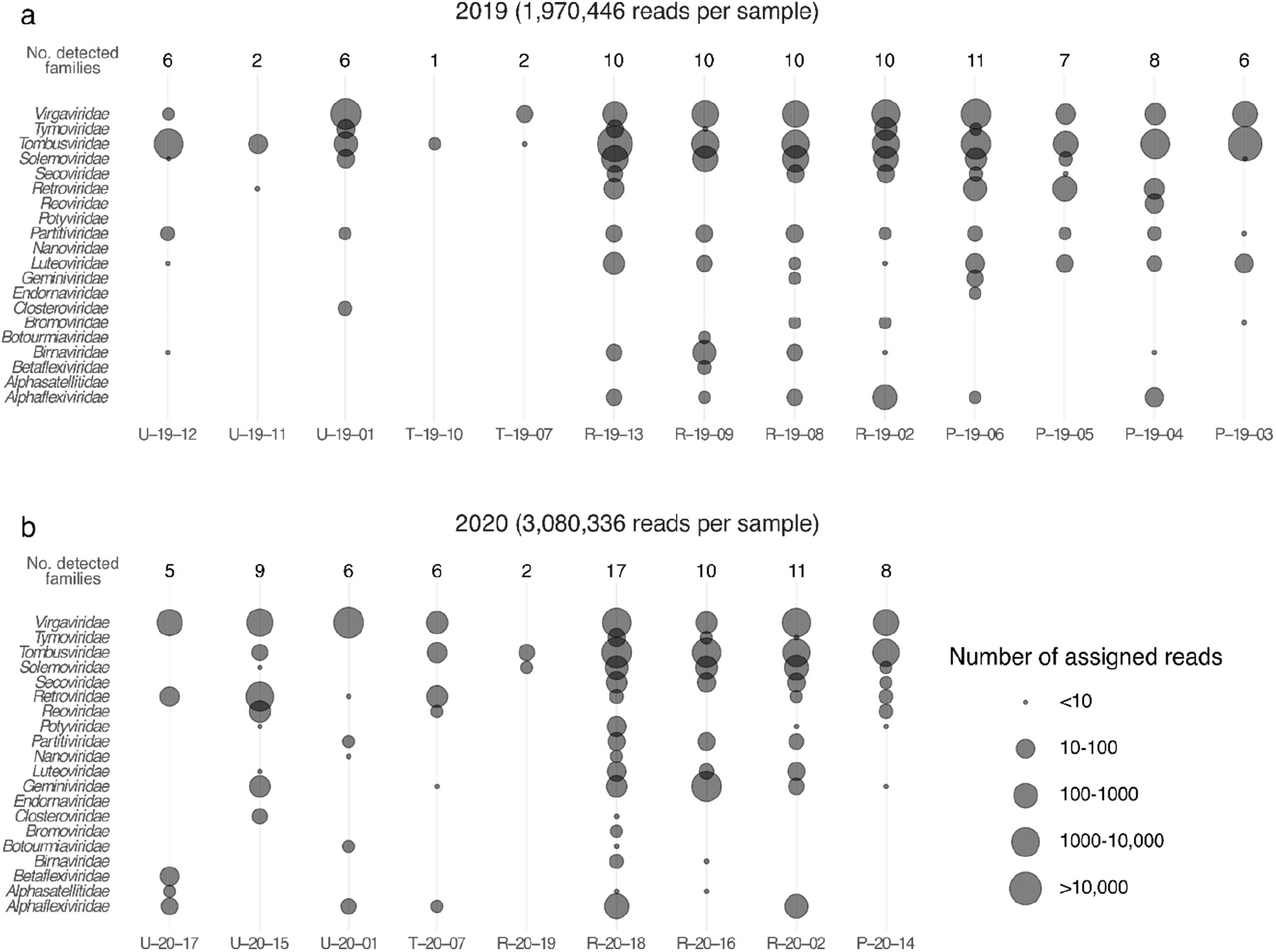
Classification of plant virus reads for different collected water samples, on the level of families. Bubble charts depict detected families of plant viruses in each sample and their corresponding read counts. The size of the bubble shows the number of assigned reads, based on DIAMOND blastx similarity search, followed by MEGAN-LCA binning. Samples are divided by sampling year: (a) 2019 and (b) 2020, due to the different subsampling size (listed next to the year designation); number of detected plant virus families for each sample is shown at the top of each panel.

Similar trend can also be corroborated by the results of the targeted qPCR tests for three selected tobamoviruses (ToMV, PMMoV, CGMMV). A difference in both the number and signal strength for tested viruses was present between the two main water subtypes (groundwater and underground water). In eight out of 13 groundwater samples, all 3 targets were detected with the strongest signal recorded for ToMV in sample P-20-14 (Cq 20). On the other hand, we did not detect all 3 viruses in any of the underground water samples and Cq values in these samples were above 25.

### 3.3. Many known plant viruses were detected in analysed water samples

We have next looked further into classification of the viral reads on the genus and species level to search for presence of known plant viruses in irrigation and other ground water samples. Reads corresponding to plant viruses from 18 different genera were detected. All of the genera were found in the groundwater samples, while only 12 of them were detected in underground water (Figure 4, 5, Supplementary Information 1, S8). Reads assigned to *Tobamovirus, Tombusvirus,* and *Sobemovirus* genera were most abundant across samples.

**Figure 4.**
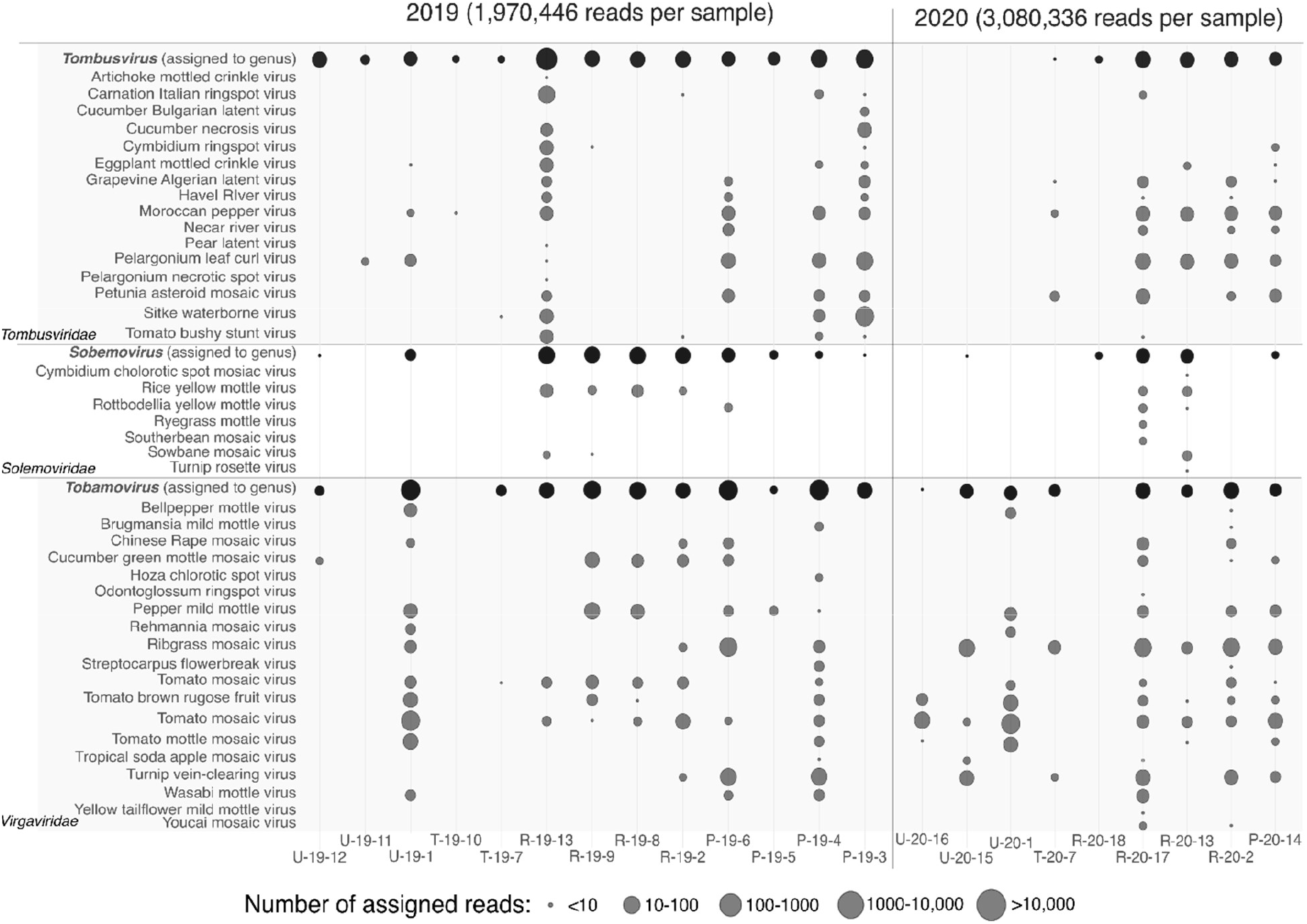
Classification of plant virus reads for three most abundant plant virus genera for different collected water samples, on the level of species and genera. Bubble charts depict detected species/genera of plant viruses in each sample and their corresponding read counts. The size of the bubble shows the number of assigned reads, based on DIAMOND blastx similarity search, followed by MEGAN-LCA binning and additional curation. Reads that could not be classified to the level of species (because they were too divergent) were classified on the level of genera (black bubbles). Samples are divided by sampling year: (a) 2019 and (b) 2020, due to the different subsampling size (listed next to the year designation).

**Figure 5.**
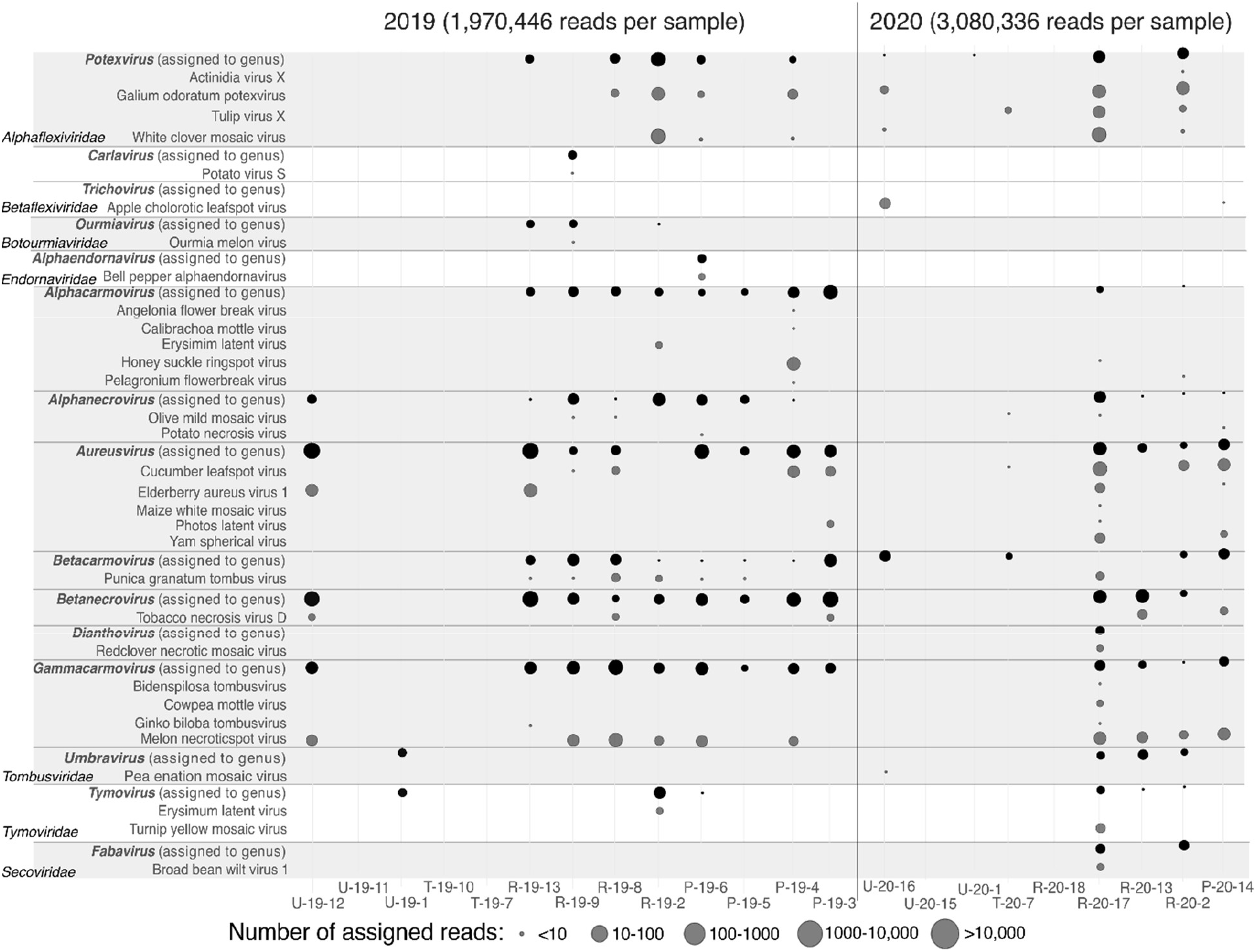
Classification of plant virus reads for fifteen less abundant plant virus genera for different collected water samples, on the level of species and genera. Bubble charts depict detected species/genera of plant viruses in each sample and their corresponding read counts. The size of the bubble shows the number of assigned reads, based on DIAMOND blastx similarity search, followed by MEGAN-LCA binning and additional curation. Reads that could not be classified to the level of species (because they were too divergent) were classified on the level of genera (black bubbles). Samples are divided by sampling year: (a) 2019 and (b) 2020, due to the different subsampling size (listed next to the year designation).

Overall, reads of 73 different plant virus species were detected (Supplementary Information 1, S8). Thirty-seven percent of detected viruses were present only in individual samples and 10 viruses had more than 100 associated reads in an individual sample (Figure 4, 5, Supplementary Information 1, S8). These viruses belong to the genus *Aureusvirus* (1 species), *Tombusvirus* (3 species) and *Tobamovirus* (6 species). Some of the species, detected in both ground and underground samples, include ToMV (detected in 14 out of 22 samples), Moroccan pepper virus (*Tombusvirus, Tombusviridae*) (11/22), tobacco mosaic virus (*Tobamovirus, Virgaviridae*) (11/10) and pelargonium leaf curl virus (*Tombusvirus, Tombusviridae*) (10/22).

Although nearly all viruses were present in ground and underground water sources, exceptions exist, such as the case of rice yellow mottle virus (*Sobemovirus, Solemoviridae*) that was detected only in groundwater, and the Rehmannia mosaic virus (*Tobamovirus, Virgaviridae*), detected only in underground water.

### 3.4. Virome analysis of water samples revealed new plant viruses present in the environment

We assembled complete or near-complete genomes of seven new viruses spanning five different genera, obtained from five different water samples (Supplementary Information 1, S11). Complete genome sequences were reconstructed for three new viruses, and partial genomic sequences, lacking one or more ORFs, were reconstructed for four new viruses.

Partial genomes of three novel viruses from three different genera were obtained: Novo mesto aureusvirus 1 (*Aureusvirus, Tombusviridae*), Gorica betanecrovirus 1 (*Betanecrovirus, Tombusviridae*), and Bericevo sobemovirus 1 (*Sobemovirus, Solemoviridae*) (Supplementary Information 3, Figure 1-3). Percent pairwise identities comparisons showed that each virus was below the species demarcation criteria for the corresponding genus (Supplementary Information 3, Figure 4-6). Based on the phylogenetic analysis, we provided details on how the new viruses clustered within their genus in Supplementary Information 3, Figure 10-12.

**Figure 6.**
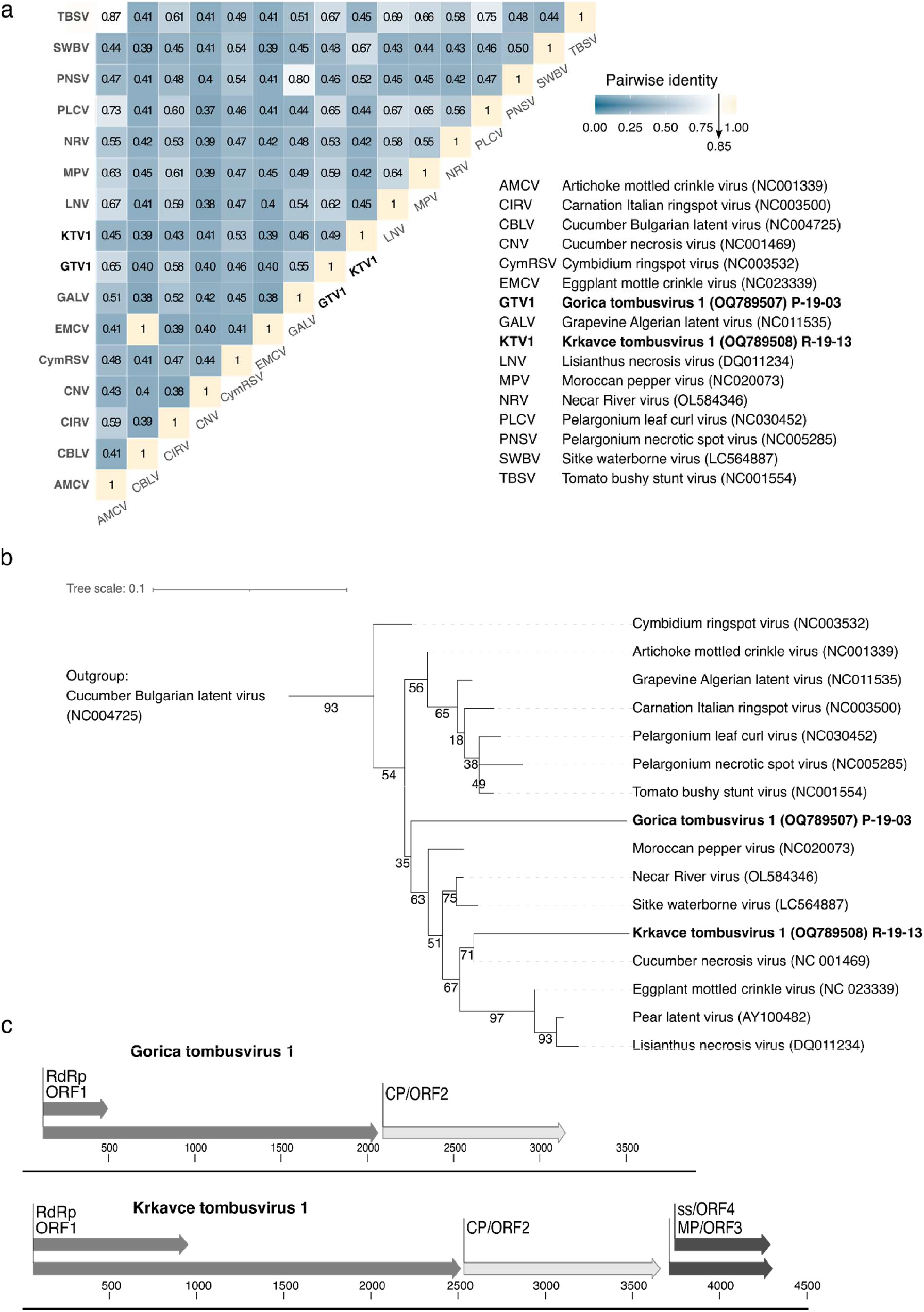
Genomic characteristics of two new tombusviruses discovered in water samples. (a) Percent pairwise identities for the amino acid sequence of coat protein (CP) for the two new viruses and members of the *Tombusvirus* genus; arrow on the colour key designate the ICTV proposed species demarcation criterion (b) Maximum likelihood phylogenetic tree based on the alignment of conserved RdRp protein sequences of the two new viruses (bolded) and members of the *Tombusvirus* genus, rooted using the outgroup (maize white line mosaic virus, *Aureusvirus*, *Tombusviridae*); numbers in brackets represent NCBI GenBank accession numbers of corresponding nucleotide sequences; numbers on the branches represent bootstrap support values; the branch length represents average number of amino acid substitutions pre site. (c) Genome structure of the two new tombusviruses with annotated predicted ORFs.

#### Tombusvirus

Partial genome sequence of Gorica tombusvirus 1 (*Tombusvirus, Tombusviridae)* (location 3, year 2019), lacking the movement protein and silencing suppressor ORFs, and complete genome sequence of Krkavce tombusvirus 1 (*Tombusvirus, Tombusviridae)* (location 13, year 2019) were reconstructed (Figure 6). Percent pairwise identities for the amino acid sequence of coat protein (CP) showed that both new viruses have 80% or less identity with other viruses in the genus, which is below the current species demarcation criteria (Figure 6). Phylogenetic analysis including tombusviruses revealed that Gorica tombusvirus 1 is not closely related to other tombusviruses, although its assignment to the genus is well supporter with bootstrap analysis, and that Krkavce tombusvirus 1 is clustering with cucumber necrosis virus (Figure 6).

#### Tobamovirus

Two complete novel tobamovirus genome sequences were reconstructed, Gorica tobamovirus 1 (*Tobamovirus, Virgaviridae*) (location 4, year 2019) and Bertoki tobamovirus 1 (*Tobamovirus, Virgaviridae*) (location 17, year 2020). Both viruses show genome composition and organisation typical of tobamoviruses (Figure 7a). Percentage pairwise identity comparisons based on complete genome nucleotide sequence revealed that both viruses have pairwise percent identities with other tobamoviruses lower than the species demarcation criteria (Supplementary Information 3, Figure 7). Phylogenetic analysis showed that Bertoki mosaic virus 1 is clustered (with high bootstrap support) in the subclade containing ribgrass mosaic virus and related viruses. Consistent with the results of the pairwise identity comparisons, Bertoki tobamovirus 1 is most closely related to PTV1. Gorica tobamovirus 1 represents more divergent branch on the tree, which still clusters with the same bigger subclade within the genus (Figure 7b).

**Figure 7.**
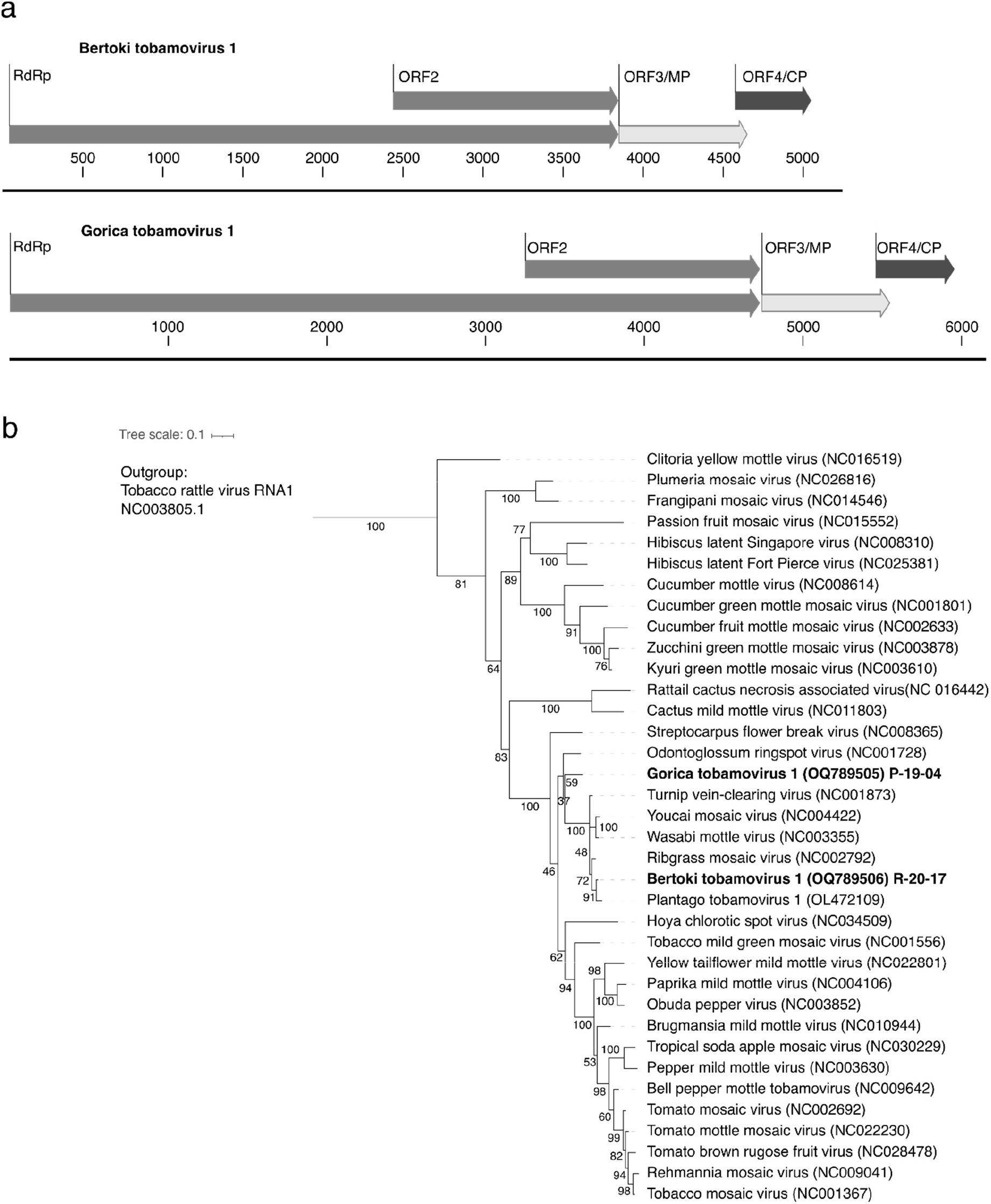
Genomic characteristics of two new tobamoviruses discovered in water samples. (a) Genome structures with annotated predicted ORFs. (b) Maximum likelihood phylogenetic tree based on the alignment of conserved RdRp protein sequences of the two new viruses (bolded) and members of the *Tobamovirus* genus, rooted using the outgroup (tobacco rattle virus, *Tobravirus*, *Virgaviridae*); numbers in brackets represent NCBI GenBank accession numbers of corresponding nucleotide sequences; numbers on the branches represent bootstrap support values; the branch length represents average number of amino acid substitutions per site.

### 3.5. Overlaps in detection of viruses in water and plant samples

In a recently published parallel study, we analysed samples of tomato and weed plants from the same locations as the water samples used in this study, to detect plant viruses [19]. Here, we now compared viruses found in plants and in water to see if there were any overlaps in detection (e.g., year, location, region). A total of 14 viruses and 2 satellite viruses were detected in both plant and water samples, disregarding the location and sampling year (Supplementary Information 1, S10). Out of those 16, four are not associated with plants as hosts, another four are new viruses detected for the first time in the previously mentioned plant virome study and the remaining 6 viruses and 2 satellite viruses are previously known to infect plants. Five plant viruses, (olive latent virus 1 (*Alphanecrovirus, Tombusviridae*), white clover mosaic virus (*Potexvirus, Alphaflexiviridae*), ToMV, and tomato bushy stunt virus (*Tombusvirus, Tombusviridae*), with tomato bushy stunt virus satellite RNA B10), were detected in both plant and water samples at the same location, at the same sampling time (Figure 8a). Instances of plant virus detection in both water and plant samples were observed in 3 out of 4 regions. However, the regional distribution of viruses detected in plants that were also detected in water varied, from detection in water in all 4 regions (i.e., ToMV), to detection in only one (i.e., olive latent virus), (Figure 8b). Additionally, also the 4 new plant viruses, which we detected and reconstructed from water for the first time in this study, were not found only in single samples; we were able to detect their nucleic acids in several different samples across the sampled sites (Supplementary Information 1, S11).

**Figure 8.**
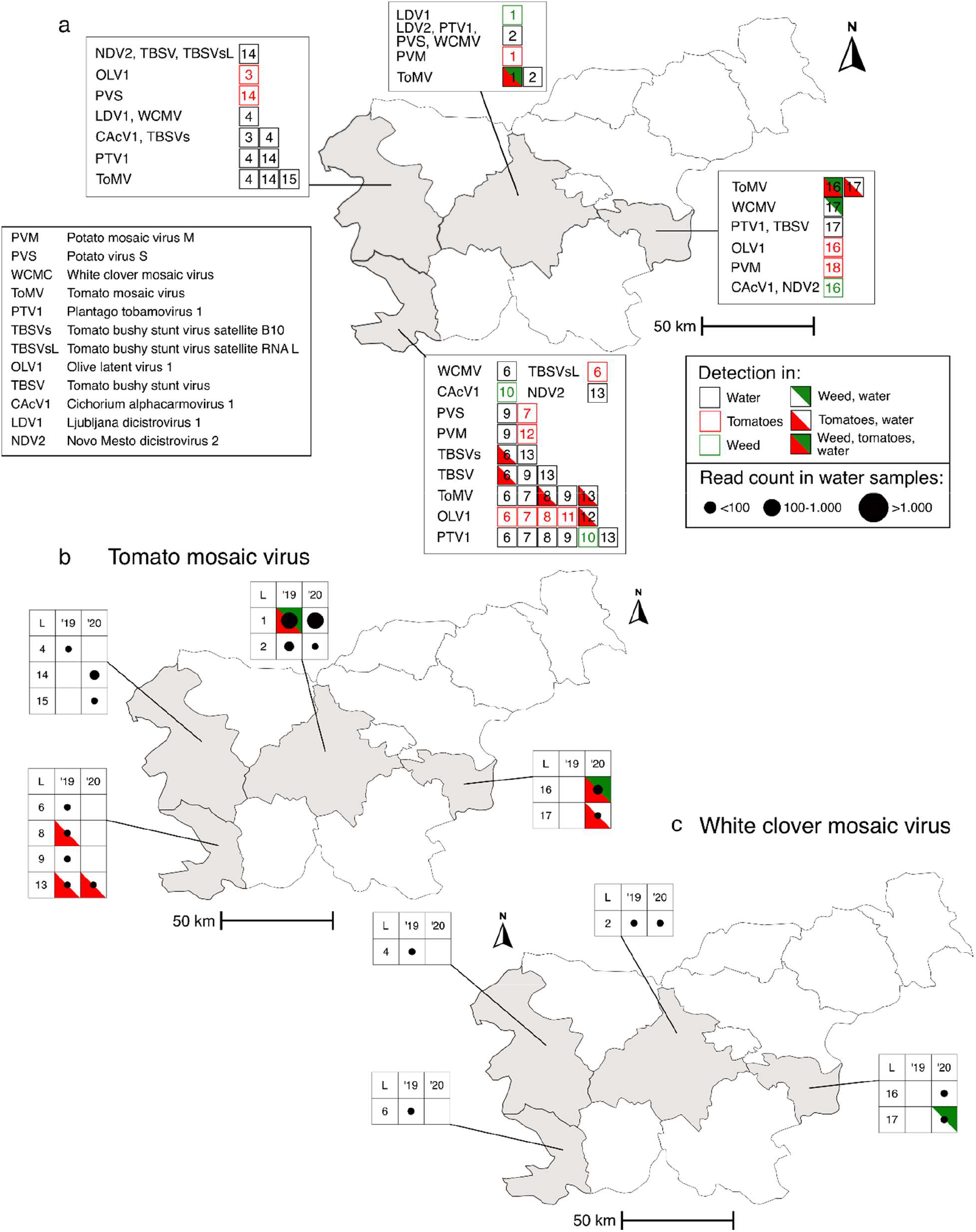
Geographical distribution of plant viruses detected both in water and plant samples. (a) Map showing the exact locations (next to the virus names’ abbreviations) of the detection of all such viruses in tomatoes, weeds, water or combination of these sample types (denoted by the colour coding explained in the legend). Viruses detected both in plants and in water at the same location were always detected in both sample types at the same sampling time. (b) – (c) Detailed results about the abundance of virus reads in water samples for two selected viruses, (b) tomato mosaic virus, (c) white clover mosaic virus. The first column of the matrices represents the location (L) where the virus was detected (Supplementary Information 1, S1), second two columns represent the years of sampling. The size of the dot in each cell corresponds to the number of virus reads in water samples. The type of the plant sample (tomato, weed) in which the virus was detected is denoted by the colour coding explained in the legend as in (a).

### 3.6. Virome analysis of water samples informs us about the diversity of a recently discovered virus

To further explore the potential of the obtained water virome data we performed a more in-depth genomic analysis of a selected plant virus, a recently discovered PTV1, which was detected in several analysed water samples. A total of eight partial consensus genomic sequences with highest similarity to this virus from seven different water sampling locations, collected across two years, were reconstructed and compared against each other and with the original virus genomic sequence reported from a plant [19]. Four of them covered the partial genomic sequence including ORF coding for RdRp and six partial genomic sequence including ORF coding for CP. Pairwise identities of sequences ranged from 89-99% (Supplementary Information 3) and phylogenetic analysis revealed clustering of isolates in subgroups (Figure 9) for both alignments. Sequence obtained from the plant clustered together with a sequence obtained from the water in the same region, in both analyses (Figure 9). Phylogenetic analysis for partial RdRp included four water-derived sequences, all from the western region of the country, and revealed two subclusters among them. Phylogenetic analysis for partial CP included six water-derived sequences and revealed several subclusters corresponding with geographic location of samples: sequences from the western part of the country clustered together and sequences from central and eastern part of the country formed a separated divergent cluster.

**Figure 9.**
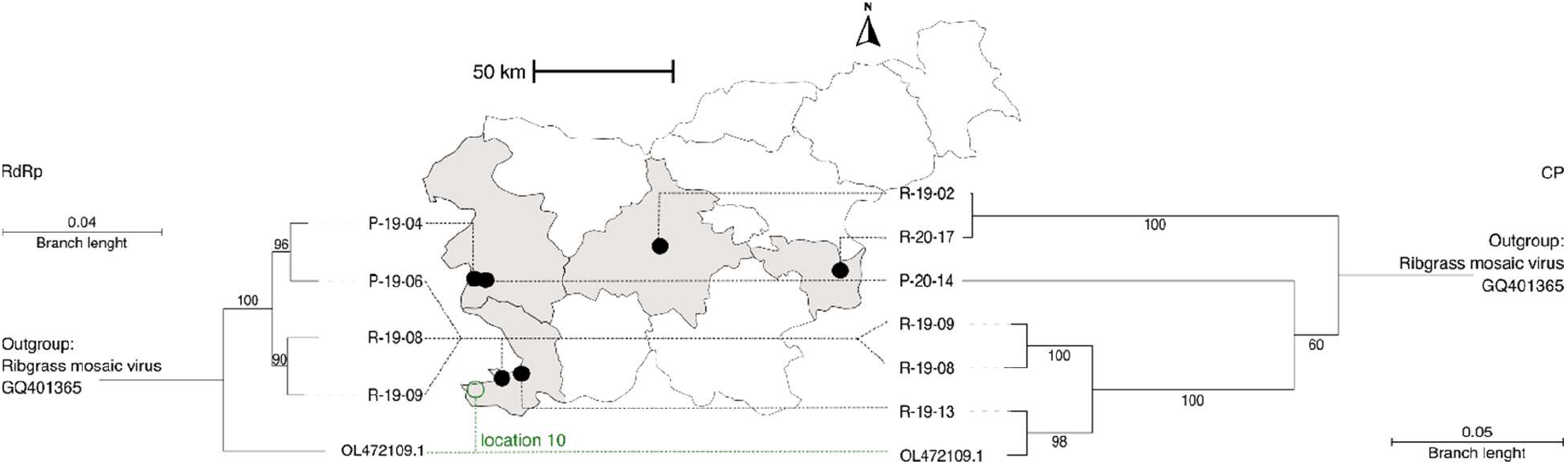
Genetic diversity of PTV1 in plant and water samples. Neighbour joining phylogenetic trees based on the alignment of partial consensus genomic sequences of PTV1 obtained from water samples and plant derived sequence from a previous study, rooted with the outgroup (ribgrass mosaic virus). Tree on the left is based on the alignment of partial genomic sequence including ORF coding for RdRp and tree on the right is based on the alignment of partial genomic sequence including ORF coding for CP. Numbers next to the branches represent bootstrap support values; the branch length represents average number of nucleotide substitutions per site. Viral genomic sequences on the tree are designated with sample numbers for water-derived consensus genomes and NCBI GenBank accession number for plant derived sample and are connected to the sampling locations as indicated on the map.

## 4. Discussion

The quality of irrigation water can play a vital role in the health and productivity of agricultural crops. Presence of plant viruses in water has been known for a long time [2], and interest in their presence in irrigation water is slowly increasing. This study described a comprehensive virome analysis that explored diversity of plant viruses in irrigation water. Shotgun HTS analysis, that was applied to water samples, shed light on the viral abundance and diversity within the samples. Viral reads accounted for around 1% of all reads, comparable to previously reported results [4], [9]. Although the sample preparation used here aimed at concentrating RNA sequences, we observed all possible variations of viral genome types within each sample. With this amount of data, we were able to detect members of 20 viral families infecting plants. There is a notable difference in both abundance and richness of detected viral families between ground and underground water sources, which can implicate diverse water quality for irrigation purposes. Two most abundantly present plant virus families in our samples, *Virgaviridae* and *Tombusviridae*, are frequently detected in aqueous environments [4], [12], [33]. Member species of both of these families have experimentally been shown to retain infectivity in extreme conditions, such as the human gastrointestinal tract [34]. Therefore, their abundant presence in water in comparison to less stable viruses is consistent with previous results. Individual species from these families are considered an economical threat to various agriculturally important plant species, mainly from *Solanaceae* family (e.g., ToMV in tomato). Out of 73 plant viruses detected in this study, ToMV was the most frequently and abundantly detected virus throughout the samples.

Using HTS-based water virome analysis we were able not only to detect wide variety of known plant viruses that circulate in the environment, but also discovered new plant viruses. Few instances of discovery of novel plant viruses in water samples exist prior the use of HTS. For example, Sikte waterborne virus, was first detected by inoculation of test plants with water from Sikte river in Germany [35]. However, it took another 15 years for its detection in plants outside the laboratory and a few more years to obtain its complete genome [35]. By analysing the sequence data obtained from water samples in this study we were able to discover seven new virus species from five different plant virus genera, reconstruct their complete or near-complete genomes, and show their presence in several water samples, often in different regions.

Within this study we were able to leverage previously available information on plant virome from the same sampling locations [19]. This enabled us to conduct a pioneering approach to compare the presence and abundance of plant viruses both in water and in plants within same agroecosystems. For example, by comparing the detection instances in water with detection in tomatoes [19], we observed a simultaneous detection of ToMV in both sample types for location 1 (Figure 8b). At the time of sampling, a large-scale outbreak of ToMV in tomato was observed in one of the greenhouses at the tomato farm that comprises this location. This brings to light the usefulness of water testing for the surveillance of plant virus outbreaks, since the viruses can be released from the plants into the environment. This concept was to some extent already explored in our previous research, where we showed that tomato brown rugose fruit virus (*Tobamovirus*, *Virgaviridae*), a damaging tomato pathogen, can be detected in water in experimental hydroponics system and in drain water from commercial greenhouses, when plants are infected [8]. As we have seen during the COVID-19 pandemic, wastewater monitoring schemes proved their applicability as an early warning system [18]. It is likely that a well-positioned (irrigation) water monitoring scheme can provide similar benefits in the context of managing plant health.

The detection of ToMV and only few other viruses in both plants and water samples from the same location and year (Figure 8a) might indicate that this pattern can be observed only if there is a high virus infection burden in the planted crops. In total, we detected 16 viruses that were previously found in plants at sampled locations [19], and also in water. However, most often they were not found in water at the same location or at the same sampling time as in plants. In many water samples, where virus reads counts were low and same viruses were not detected in nearby plants, deducing the connection between the plants and water is not possible. Nevertheless, detection of plant viruses in water can have several important implications. For example, the detection of important plant pathogen (e.g., a virus on quarantine list for a region) in water samples can motivate increased survey for the virus in nearby host plants or intensified research on the characteristics of such virus in the environment [8]. Moreover, as we have demonstrated in this study, water virome data can rapidly provide a wealth of information about possible distribution of known and newly discovered viruses in the environment. For example, white clover mosaic virus was detected in plant virome analysis in a single plant sample on studied locations, however, addition of data from water analysis indicated much wider distribution of virus in the environment, since it was detected in nearly all water samples, covering all analyzed regions (Figure 8c). However, this study also highlights the limitations of using water samples as a proxy for distribution of plant viruses, as the lack of detection in water does not guarantee lack of detection in plants, like in the example of olive latent virus 1, which was detected at seven different locations in plants, but at only one in water (Figure 8a). These discrepancies can be expected, since many different factors such as virus titer in plants, abundance and distributions of host plants, and virus stability in the environment, would affect the possible presence, abundance and spread of different viruses in environment.

HTS-based virome studies result in large sequence datasets that can be further exploited to study not only presence, but also diversity of viruses in a region under study. In this study, we demonstrated that water virome data can bring extensive additional insights into diversity of recently discovered viruses, for which limited information is available. Plantago tobamovirus 1 was recently discovered in a single plant sample [19] at one of our sampling locations. Here, we detected it in multiple water samples, covering all regions under study. We reconstructed partial consensus genome sequences of this virus from several water samples and confirmed notable genetic diversity within the species. Moreover, clustering of isolates derived from water virome data is largely matching the geographic distribution of sampled waters. Thus, having only a single occurrence and genomic information about the virus from its plant host, using water virome data, we were able to discover that virus is likely widespread and genetically diversified in the studied ecosystems. Interestingly, closely related tobamovirus from the ribgrass mosaic virus subgroup have been isolated from river water from Hungary in 1986 [36]. Together with results from this study, this opens interesting questions about the wide presence and survival of these viruses and abundance of their hosts in ecosystems, likely close to the surface water bodies.

Within the irrigation water virome study presented here, we obtained rich information about the presence of plant viruses in the region (also for some that were previously not reported in the country). We demonstrated usability of environmental water virome analysis as a tool for studying the diversity and ecology of plant viruses, as well as for the discovery of novel virus species. In contrast to more traditional plant virus surveillance targeting host plants, water virome analysis allows monitoring of a larger area with a single sample and might provide early warning prior to large outbreaks in agroecosystems. Additional research should be focused on assessing the biological relevance of new viruses found in environmental irrigation waters and to study the directionality of spread between viruses found in water and plant components of the ecosystem. Moreover, HTS-based analysis of irrigation waters can be employed to study within-species diversity of plant viruses and detect potentially emergent virus variants that could represent threat for global plant health.

## Conclusions

- HTS-based virome analysis of irrigation and other ground water sources informed us about the presence of a wide range of plant viruses in the regions under study, with environmentally stable viruses, such as tobamoviruses and tombusviruses, being the most abundantly detected in the analysed water sources.
- Notable differences in both abundance and richness of plant virus nucleic acids were observed between different water types, whereas ground water samples in general had higher count of plant viruses present in higher abundance, which might imply different quality of investigated water types.
- The information obtained from water virome analysis allowed us to detect both known viruses previously not found in an area and new virus species, which overall, provided us with a clearer understanding of environmental presence of a given virus and its diversity in the agroecosystem.
- Understanding the presence and diversity of plant viruses in irrigation water might be a contributing factor in ensuring effective management of plant health in future. Water virome analysis was shown here to be a useful tool for the surveillance and discovery of known and new plant viruses in a wider environment.

## Data availability

HTS data produced in this study is available via European Nucleotide Archive (ENA) under project code PRJEB60028. Complete or partial genomes of new virus species discovered in this study were submitted to NCBI GenBank, and their accession numbers can be found in Supplementary Information 1, S11. Consensus genomic sequences of PTV1 obtained from water samples are available in Supplementary Information 2. Codes used for additional reads classification curation and generation of figures in R are available in Supplementary Information 2.

### Abbreviations

RT-qPCR: reverse transcription quantitative polymerase chain reaction
HTS: high throughput sequencing
PMMoV: pepper mild mottle virus
ToMV: tomato mosaic virus
CGMMV: cucumber green mottle mosaic virus
LUC: Luciferase Control RNA
NCI: negative control of isolation
CLC-GWB: CLC Genomics Workbench
LCA: lowest common ancestor
ICTV: International Committee on Taxonomy of Viruses
ORF: open reading frame
(-) ssRNA: negative sense single stranded RNA
(+) ssRNA: positive sense single stranded RNA
dsRNA: double stranded RNA
PTV1: plantago tobamovirus 1
RdRp: RNA-dependent RNA polymerase

## Acknowledgements

We would like to thank all the farm owners for the access to and permission for water sampling and Mr Gabrijel Seljak for his help in organisation and communication with farm owners. This work was supported by the funding from Horizon 2020 Marie Skłodowska-Curie Actions Innovative Training Network (H2020 MSCA-ITN) project “INEXTVIR” (GA 813542) under the management of the European Commission-Research Executive Agency; from the Slovenian Research and Innovation Agency (ARIS) core financing P4-0407 and P4-0165 and applied research project L4-3179 (Discovery and water-linked epidemiology of emergent tobamoviruses infecting crops) funded by ARIS, the Ministry of Agriculture and Food of the Republic of Slovenia and BIA d. o. o. Laboratory and process equipment company (Ljubljana, Slovenia).

## Notes

### Competing Interest Statement

This work was supported by the funding from Horizon 2020 Marie Skodowska-Curie Actions Innovative Training Network (H2020 MSCA-ITN) project INEXTVIR (GA 813542) under the management of the European Commission-Research Executive Agency; from the Slovenian Research and Innovation Agency (ARIS) core financing P4-0407 and P4-0165 and applied research project L4-3179 (Discovery and water-linked epidemiology of emergent tobamoviruses infecting crops) funded by ARIS, the Ministry of Agriculture and Food of the Republic of Slovenia and BIA d. o. o. Laboratory and process equipment company (Ljubljana, Slovenia).

## Bibliography

[1] K. S. Sastry and T. A. Zitter, “Management of Virus and Viroid Diseases of Crops in the Tropics,” in Plant Virus and Viroid Diseases in the Tropics: Volume 2: Epidemiology and Management, K. S. Sastry and T. A. Zitter, Eds., Dordrecht: Springer Netherlands, 2014, pp. 149–480. doi: 10.1007/978-94-007-7820-7_2.

[2] R. Koenig, “Plant viruses in rivers and lakes,” Advances in Virus Research, vol. 31, no. C, pp. 321–333, 1986, doi: 10.1016/S0065-3527(08)60267-5.

[3] N. Mehle and M. Ravnikar, “Plant viruses in aqueous environment - Survival, water mediated transmission and detection,” Water Research, vol. 46, no. 16, pp. 4902–4917, 2012, doi: 10.1016/j.watres.2012.07.027.

[4] K. Bačnik et al., “Viromics and infectivity analysis reveal the release of infective plant viruses from wastewater into the environment,” Water Research, pp. 115628–115628, Feb. 2020, doi: 10.1016/j.watres.2020.115628.

[5] K. Rosario, C. Nilsson, Y. W. Lim, Y. Ruan, and M. Breitbart, “Metagenomic analysis of viruses in reclaimed water,” Environmental Microbiology, vol. 11, no. 11, pp. 2806–2820, 2009, doi: 10.1111/j.1462-2920.2009.01964.x.

[6] J. Chopyk et al., “Comparative metagenomic analysis of microbial taxonomic and functional variations in untreated surface and reclaimed waters used in irrigation applications,” Water Research, vol. 169, p. 115250, Feb. 2020, doi: 10.1016/j.watres.2019.115250.

[7] J. A. Rothman and K. L. Whiteson, “Sequencing and Variant Detection of Eight Abundant Plant-Infecting Tobamoviruses across Southern California Wastewater,” Microbiology Spectrum, vol. 10, no. 6, pp. e03050-22, Nov. 2022, doi: 10.1128/spectrum.03050-22.

[8] N. Mehle et al., “Tomato brown rugose fruit virus in aqueous environments – survival and significance of water-mediated transmission,” Frontiers in Plant Science, 2023.

[9] A. Lopez-Roblero, D. J. Martínez Cano, E. Diego-García, G. K. Guillén-Navarro, P. Iša, and E. Zarza, “Metagenomic analysis of plant viruses in tropical fresh and wastewater,” Environmental DNA, vol. n/a, no. n/a, doi: 10.1002/edn3.416.

[10] S. Vani and A. Varma, “Properties of cucumber green mottle mosaic virus isolated from water of river Jamuna.,” Indian Phytopathology, vol. 46, no. 2, pp. 118–122, 1993.

[11] J. Boben et al., “Detection and quantification of Tomato mosaic virus in irrigation waters,” Eur J Plant Pathol, vol. 118, no. 1, pp. 59–71, May 2007, doi: 10.1007/s10658-007-9112-1.

[12] X. Zhang et al., “Potential risk of plant viruses entering disease cycle in surface water in protected vegetable growing areas of Eastern China,” PLOS ONE, vol. 18, no. 1, p. e0280303, Jan. 2023, doi: 10.1371/journal.pone.0280303.

[13] R. Rossi, “Irrigation in EU agriculture.” European Parlimentary Research Service, EPRS, Dec. 2019. [Online]. Available: https://www.europarl.europa.eu/RegData/etudes/BRIE/2019/644216/EPRS_BRI(2019)64421 6_EN.pdf

[14] F. H. Coutinho et al., “Marine viruses discovered via metagenomics shed light on viral strategies throughout the oceans,” Nat Commun, vol. 8, no. 1, Art. no. 1, Jul. 2017, doi: 10.1038/ncomms15955.

[15] C. Gao et al., “Virioplankton assemblages from challenger deep, the deepest place in the oceans,” iScience, vol. 25, no. 8, p. 104680, Aug. 2022, doi: 10.1016/j.isci.2022.104680.

[16] J. Lu et al., “Metagenomic analysis of viral community in the Yangtze River expands known eukaryotic and prokaryotic virus diversity in freshwater,” Virologica Sinica, vol. 37, no. 1, pp. 60–69, Feb. 2022, doi: 10.1016/j.virs.2022.01.003.

[17] N. Mehle, I. Gutiérrez-Aguirre, D. Kutnjak, and M. Ravnikar, “Water-Mediated Transmission of Plant, Animal, and Human Viruses,” Advances in Virus Research, vol. 101, 2018, doi: 10.1016/bs.aivir.2018.02.004.

[18] S. Shah, S. X. W. Gwee, J. Q. X. Ng, N. Lau, J. Koh, and J. Pang, “Wastewater surveillance to infer COVID-19 transmission: A systematic review,” Science of The Total Environment, vol. 804, p. 150060, Jan. 2022, doi: 10.1016/j.scitotenv.2021.150060.

[19] M. P. S. Rivarez et al., “In-depth study of tomato and weed viromes reveals undiscovered plant virus diversity in an agroecosystem,” Microbiome, vol. 11, no. 1, p. 60, Mar. 2023, doi: 10.1186/s40168-023-01500-6.

[20] E. Haramoto et al., “Occurrence of Pepper Mild Mottle Virus in Drinking Water Sources in Japan,” Appl Environ Microbiol, vol. 79, no. 23, pp. 7413–7418, Dec. 2013, doi: 10.1128/AEM.02354-13.

[21] X. Zhao et al., “Comparison of three magnetic-bead-based RNA extraction methods for detection of cucumber green mottle mosaic virus by real-time RT-PCR,” Arch Virol, vol. 160, no. 7, pp. 1791–1796, Jul. 2015, doi: 10.1007/s00705-015-2444-9.

[22] N. Toplak, V. Okršlar, D. Stanič-Racman, K. Gruden, and J. Žel, “A high-throughput method for quantifying transgene expression in transformed plants with real-time PCR analysis,” Plant Mol Biol Rep, vol. 22, no. 3, pp. 237–250, Sep. 2004, doi: 10.1007/BF02773134.

[23] B. Buchfink, C. Xie, and D. H. Huson, “Fast and sensitive protein alignment using DIAMOND,” Nature Methods, vol. 12, no. 1, pp. 59–60, 2014, doi: 10.1038/nmeth.3176.

[24] D. H. Huson et al., “MEGAN Community Edition - Interactive Exploration and Analysis of Large-Scale Microbiome Sequencing Data,” PLOS Computational Biology, vol. 12, no. 6, p. e1004957, Jun. 2016, doi: 10.1371/journal.pcbi.1004957.

[25] “ICTV Report Chapters | ICTV,” May 2022. https://ictv.global/report

[26] A. Bankevich et al., “SPAdes: A new genome assembly algorithm and its applications to single-cell sequencing,” Journal of Computational Biology, vol. 19, no. 5, pp. 455–477, 2012, doi: 10.1089/cmb.2012.0021.

[27] S. Kumar, G. Stecher, M. Li, C. Knyaz, and K. Tamura, “MEGA X: Molecular Evolutionary Genetics Analysis across Computing Platforms,” Mol Biol Evol, vol. 35, no. 6, pp. 1547–1549, Jun. 2018, doi: 10.1093/molbev/msy096.

[28] B. M. Muhire, A. Varsani, and D. P. Martin, “SDT: A Virus Classification Tool Based on Pairwise Sequence Alignment and Identity Calculation,” PLOS ONE, vol. 9, no. 9, p. e108277, Sep. 2014, doi: 10.1371/journal.pone.0108277.

[29] S. Capella-Gutiérrez, J. M. Silla-Martínez, and T. Gabaldón, “trimAl: a tool for automated alignment trimming in large-scale phylogenetic analyses,” Bioinformatics, vol. 25, no. 15, pp. 1972–1973, Aug. 2009, doi: 10.1093/bioinformatics/btp348.

[30] J. Trifinopoulos, L.-T. Nguyen, A. von Haeseler, and B. Q. Minh, “W-IQ-TREE: a fast online phylogenetic tool for maximum likelihood analysis,” Nucleic Acids Research, vol. 44, no. W1, pp. W232–W235, Jul. 2016, doi: 10.1093/nar/gkw256.

[31] I. Letunic and P. Bork, “Interactive Tree Of Life (iTOL) v5: an online tool for phylogenetic tree display and annotation,” Nucleic Acids Research, vol. 49, no. W1, pp. W293–W296, Jul. 2021, doi: 10.1093/nar/gkab301.

[32] P. I. Bustamante and R. Hull, “Plant virus gene expression strategies,” Electronic Journal of Biotechnology, vol. 1, no. 2, pp. 13–24, Aug. 1998, doi: 10.4067/S0717-34581998000200003.

[33] M. F. Duarte et al., “Metagenomic analyses of plant virus sequences in sewage water for plant viruses monitoring,” Trop. plant pathol., Apr. 2023, doi: 10.1007/s40858-023-00575-8.

[34] T. Zhang et al., “RNA Viral Community in Human Feces: Prevalence of Plant Pathogenic Viruses,” PLOS Biology, vol. 4, no. 1, p. e3, Dec. 2005, doi: 10.1371/journal.pbio.0040003.

[35] T. Uehara-Ichiki et al., “Complete genome sequence of Sikte (Sitke) waterborne virus, a member of the genus Tombusvirus,” Arch Virol, vol. 166, no. 3, pp. 991–994, Mar. 2021, doi: 10.1007/s00705-020-04949-0.

[36] N. Juretic, . Horvath, and M. Krajacic, “Occurrence of a tobamovirus similar to ribgrass mosaic virus in water of Hungarian river Zala,” Acta Phytopathologica et Entomologica Hungarica, vol. 21, pp. 291–295, 1986.

