## Supplementary Information 2 for "Virome analysis of irrigation water sources provides extensive insights into the diversity and distribution of plant viruses in agroecosystems": scripts_for_graphs.R

```
# ALL graphs

#Domains of life and genome organisation
library(ggplot2)
df_s1 <- read.table("~/domains.csv", header=TRUE, sep=";", dec=",")

ggplot(subset(df_s1, Type %in% c("Unclassified", "Evnironmental sequences", "Unclassified RNA", "ssDNA", "dsRNA", "dsDNA", "(+)ssRNA", "(-)ssRNA")), aes(x=Perc, y=SampleID, fill=Type)) +
  geom_col(position="stack", width=0.8, alpha=0.9) + geom_text(aes(label=Perc), position = position_stack(vjust=0.5, reverse=FALSE), hjust=0.5, size=3) +
  theme_minimal() + theme(panel.grid.major.x = element_blank(), panel.grid.minor.x = element_blank(), axis.title = element_blank(), legend.position = "bottom") + 
  facet_wrap(vars(Year), nrow = 2, ncol = 1, scales = "free_y") + 
  scale_fill_manual(name="Genome structure", values = c("grey45", "red", "grey55", "grey25", "grey35", "grey75", "grey10", "grey65"))  

ggplot(subset(df_s1, Type %in% c("Bacteria", "Archaea", "Eukaryota", "Viruses", "Other", "Unclassified reads")), aes(x=Perc, y=SampleID, fill=Type)) +
  geom_col(position="stack", width=0.8, alpha=0.9) + geom_text(aes(label=Perc), colour="red", position = position_stack(vjust=0.5, reverse=FALSE), hjust=0.5, size=3) +
  theme_minimal() + theme(panel.grid.major.x = element_blank(), panel.grid.minor.x = element_blank(), axis.title = element_blank(), legend.position = "bottom") + 
  facet_wrap(vars(Year), nrow = 2, ncol = 1, scales = "free_y") + 
  scale_fill_manual(name="Realms", values = c("grey15", "grey50", "grey35", "grey55", "red"))


#family
library(ggplot2)
heatmaps_family_uncl <- read.csv("~\\heatmap_family_long_no_unclas.csv", sep=";")

ggplot(heatmaps_family_uncl, aes(x=SampleID, y=Family)) +
  geom_point(aes(size=LOG10), alpha=0.5) + scale_size(range = c(1, 10)) +
  facet_wrap(vars(Year), nrow = 2, ncol = 1, scales = "free") +
  theme_minimal() + theme(panel.grid.major.y = element_blank(), legend.position = "bottom")

#host
library(ggplot2)
heatmaps_host <- read.csv("~\\heatmaps_host.csv", sep=";")

ggplot(heatmaps_host, aes(Y, Z, fill=X)) +
  geom_col(position="dodge") +
  geom_text(aes(label=Z, group=X), position=position_dodge2(width=0.8), vjust=-0.2, hjust=0.5) +
  facet_wrap(vars(Year), nrow=2, ncol=1, scales = "free") +
  theme_minimal() + theme(panel.grid = element_blank(), axis.line.x.bottom = element_line(), axis.ticks.x = element_line(), legend.position = "bottom") +
  scale_fill_manual(name="Host", values = c("grey35", "grey55", "red", "grey25")) + 
  coord_cartesian(clip="off")


#genus and species counts
library(ggplot2)
heatmaps_species <- read.csv("~\\heatmap_species_long.csv", sep=";")

ggplot(heatmaps_species, aes(x=SampleID, y=fct_rev(fct_inorder(Virus)))) +
  geom_point(aes(size=LOG10), alpha=0.5) + scale_size(range=c(1, 10)) +
  facet_grid(cols=vars(Year), scales="free_x", space="free_x") +
  theme_minimal() + theme(panel.grid.major.y = element_blank(), legend.position = "bottom") + 
  coord_cartesian(clip="off")

#minimaps
library("ggpubr")
library("ggplot2")
library("sf")
library("rnaturalearth")
library("rnaturalearthdata")
library("rnaturalearthhires")

world <- ne_countries(scale = "large", returnclass = "sf")
class(world)

locations <- read.table("~/virus_maps_all.txt", 
                        header=TRUE, sep="\t", dec=",", colClasses = c("Year"="character"))

#tomato mosaic virus
ToMV <- ggplot(data = world) +
  geom_sf() +
  geom_point(data = subset(locations, Virus %in% c("Tomatomosaicvirus")), aes(x = Longitude, y = Latitude, color=Year, size=ReadCount)) +
  coord_sf(xlim = c(13.3, 16.5), ylim = c(45.4, 46.9), expand = TRUE) +
  scale_size(limits = c(1,6000), breaks = c(1, 10, 100, 1000, 6000), labels=c("1-10", "10-100", "100-1000", "1000-5000", ">5000"), range = c(2,12)) +
  theme_void()
ToMV

#Tomato bushy stunt virus
TBSV <- ggplot(data = world) +
  geom_sf() +
  geom_point(data = subset(locations, Virus %in% c("Tomatobushystuntvirus")), aes(x = Longitude, y = Latitude, color=Year, size=ReadCount)) +
  coord_sf(xlim = c(13.3, 16.5), ylim = c(45.4, 46.9), expand = TRUE) +
  scale_size(limits = c(1,6000), breaks = c(1, 10, 100, 1000, 6000), labels=c("1-10", "10-100", "100-1000", "1000-5000", ">5000"), range = c(2,12))
TBSV

#plantago tobamovirus 1
nV2 <- ggplot(data = world) +
  geom_sf() +
  geom_point(data = subset(locations, Virus %in% c("newVirga2")), aes(x = Longitude, y = Latitude, color=Year, size=ReadCount)) +
  coord_sf(xlim = c(13.3, 16.5), ylim = c(45.4, 46.9), expand = TRUE) +
  scale_size(limits = c(1,6000), breaks = c(1, 10, 100, 1000, 6000), labels=c("1-10", "10-100", "100-1000", "1000-5000", ">5000"), range = c(2,12)) +
  theme(axis.title.x = element_blank(), axis.title.y = element_blank())
nV2

#RT-qPCR
library(ggplot2)

qPCR <- read.csv("~\\qPCR1.csv", sep=";")

ggplot(qPCR) +
  geom_segment( aes(x=SampleID, xend=SampleID, y=raw, yend=elution), color="grey", size=1) +
  geom_point( aes(x=SampleID, y=raw), size=2 ) +
  geom_point( aes(x=SampleID, y=elution), alpha=0.65, size=5 ) +
  facet_grid(cols = vars(Year), rows = vars(Type), scales = "free", space="free") +
  theme_minimal() + theme(panel.grid.major.y = element_blank(), panel.grid.minor.y = element_blank()) +
  labs(x="Sample ID", y="Cq value")
```
