## Supplementary Information 2 for "Virome analysis of irrigation water sources provides extensive insights into the diversity and distribution of plant viruses in agroecosystems": generic_names.txt

#!/bin/bash

# navigate to the location where megan parser is installed in
cd $path/megan6_parser/

# run megan parser on a specific .blast table; species demarcation criteria is given as an argument of the script 
mix run ./tasks/parse.exs $path/balst_table.blast $1 $path/filename_parsed.csv

#return to the location where the data is stored
cd $path

#remove first like of the .csv file
sed 1d filename_parsed.csv > filename_parsed_ready.csv

#extract only accesion numbers
csvtool col 2 filename_parsed_ready.csv > filename_accno.txt

#using EntrezTools retrieve generic name for each accession number 
for LINE in `cat filename_accno.txt`
do
   esearch -db protein -query $LINE | efetch -format docsum | xtract -pattern DocumentSummary -element Organism >> filename_generic.txt
done

#check if all generic names are retreived 
if [ "$(wc -l < filename_accno.txt)" -eq "$(wc -l < filename_generic.txt)" ] 
then 
	echo 'Match!'

# create the db of accession numbers and corresponding generic names
	csvtool pastecol 2 1 filenameaccno.txt filename_generic.txt > filename_db.csv
# sorting the db and pared file based on the id column and matching the virus name
	csvtool pastecol 5 2 filename_parsed_ready.csv filename_db.csv > filename_final.csv
# sorting the final file based on the read id and score
	sort -k1 -k3 filename_final.csv > filename_final_sorted.csv
# taking only one line per read with the highest score
	awk -F, 'k&&k!=$1{print p}{p=$0;k=$1}END{print p}' filename_final_sorted.csv > filename_DONE.csv

else 
	echo 'Warning: No Match!'
fi
