## Supplementary Information 2 for "Virome analysis of irrigation water sources provides extensive insights into the diversity and distribution of plant viruses in agroecosystems": README.md

# Megan6Parser

This is a small parser to read MEGAN6 output files.
After obtaining a meaningful structure, work is being done to manipulate and generate desired outputs.

## Installation

```elixir
mix deps.get
```

## Running the parser

```elixir
mix run ./tasks/parse.exs <input_file::string> <criteria::number> <output_file::string> <db_file::optional_string>
```

Not providing the `<db_file::optional_string>` parameter will do a search without a database, which will not try to match the English names.
This is far more efficient.

Documentation can be generated with [ExDoc](https://github.com/elixir-lang/ex_doc)
and published on [HexDocs](https://hexdocs.pm). Once published, the docs can
be found at [https://hexdocs.pm/oli_parser](https://hexdocs.pm/oli_parser).
