## Supplementary Information 2 for "Virome analysis of irrigation water sources provides extensive insights into the diversity and distribution of plant viruses in agroecosystems": readme.docx

Supplementary Information 2 contains following files/folders:

- figures: .csv files related to each figure in the paper and an R script use to generate raw figures. All figures were later edited in Inkscape
- PTV1_ alignment_RdRp/CP: .fasta files containing the alingment of consensus sequences used in plantago tobamovirus 1 analysis for each of the two sections
- megan6_parser-master: The script takes as input a BLAST table of a given genus and a numeric value indicating the species demarcation criteria for the given genus (retrieved from ICTV). The return is a CSV table with Read ID, NCBI accession number, percent identity, and status. The first three columns are information from the original BLAST table and the script adds the fourth column. It has a binary output, YES or NO, indicating whether the percent identity for a given match is above or below the specified species demarcation criteria. In addition to assigning the status column, the script also reduces the table BLAST, keeping only the best match for each read. Finally, generic names of the viruses are determined using a set of tools available from Entrez Direct (Kans, 2022).
- generic_names: bash script that deals with assigning generic names is given separately.

[Kans, J. (2022). Entrez Direct: E-utilities on the Unix Command Line. In *Entrez Programming Utilities Help [Internet]*. National Center for Biotechnology Information (US). https://www.ncbi.nlm.nih.gov/books/NBK179288/](Kans,%20J.%20(2022).%20Entrez%20Direct:%20E-utilities%20on%20the%20Unix%20Command%20Line.%20In%20Entrez%20Programming%20Utilities%20Help%20%5bInternet%5d.%20National%20Center%20for%20Biotechnology%20Information%20(US).%20https://www.ncbi.nlm.nih.gov/books/NBK179288/)
