## Supplementray Information 3 for "Virome analysis of irrigation water sources provides extensive insights into the diversity and distribution of plant viruses in agroecosystems"

### Supplementary Information 3

Supplementary figure 1-3 - Genome organisation of novel viruses showingg open reading frames and the poteins it codes for. Genome length in number of bases are shown on scale. For GenBank accession numbers refer to upplementary Information 1.

#### Supplementary Figure 1 - Novo mesto aureusvirus 1

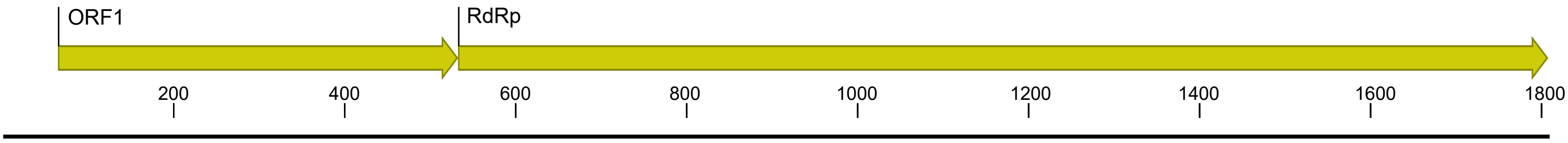


#### Supplementary Figure 2 - Gorica betanecrovirus 1

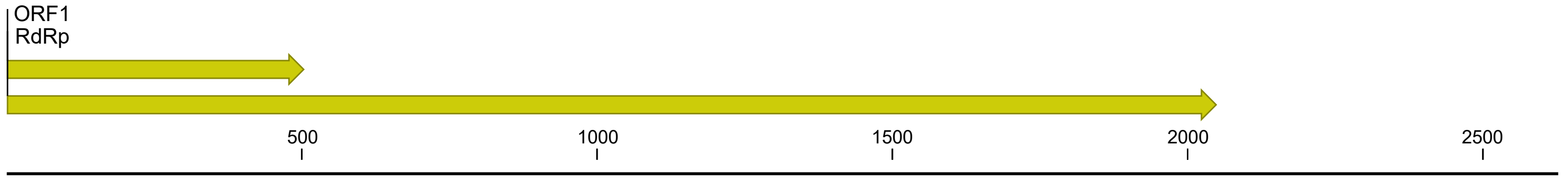


#### Supplementary Figure 3 - Bericevo sobemovirus 1

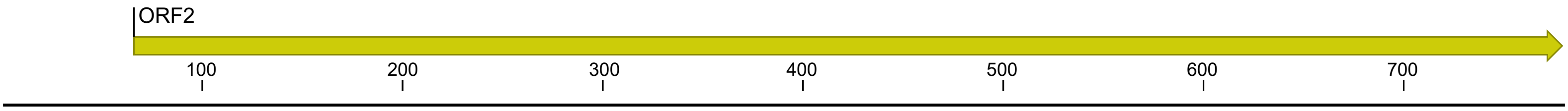


Supplementary figure 4-7 - Pairwise similarity comparison based on the species demarcation criteria as listed in ICTV (details in Supplementary Information 1). GenBank accession numbers are listed next to known sequences. For GenBank accesion numbers of novel viruses refer to Supplementary Information 1.

#### Supplementary Figure 4 - Pairwise similariy comparison of virus species from *Betanecrovirus*

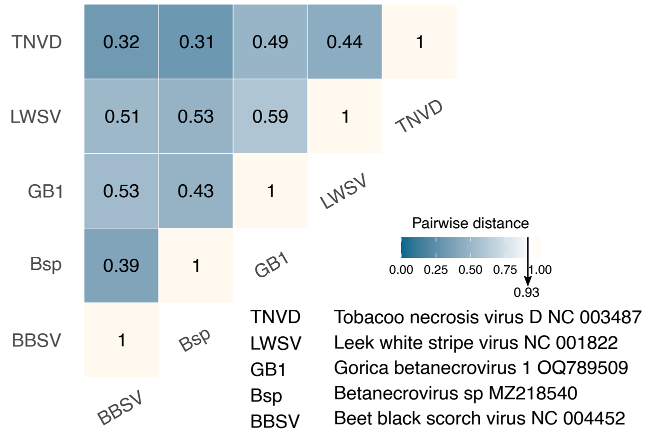

#### Supplementary Figure 5 - Pairwise similariy comparison of virus species from *Aureusvirus*

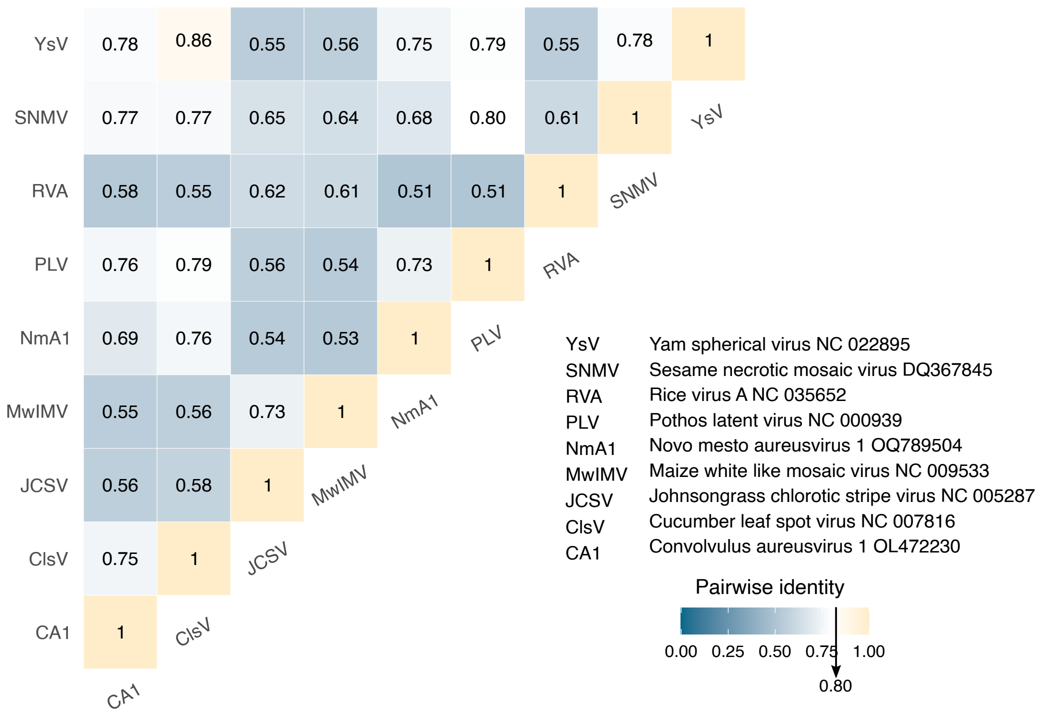


#### Supplementary Figure 6 - Pairwise similariy comparison of virus species from *Sobemovirus*

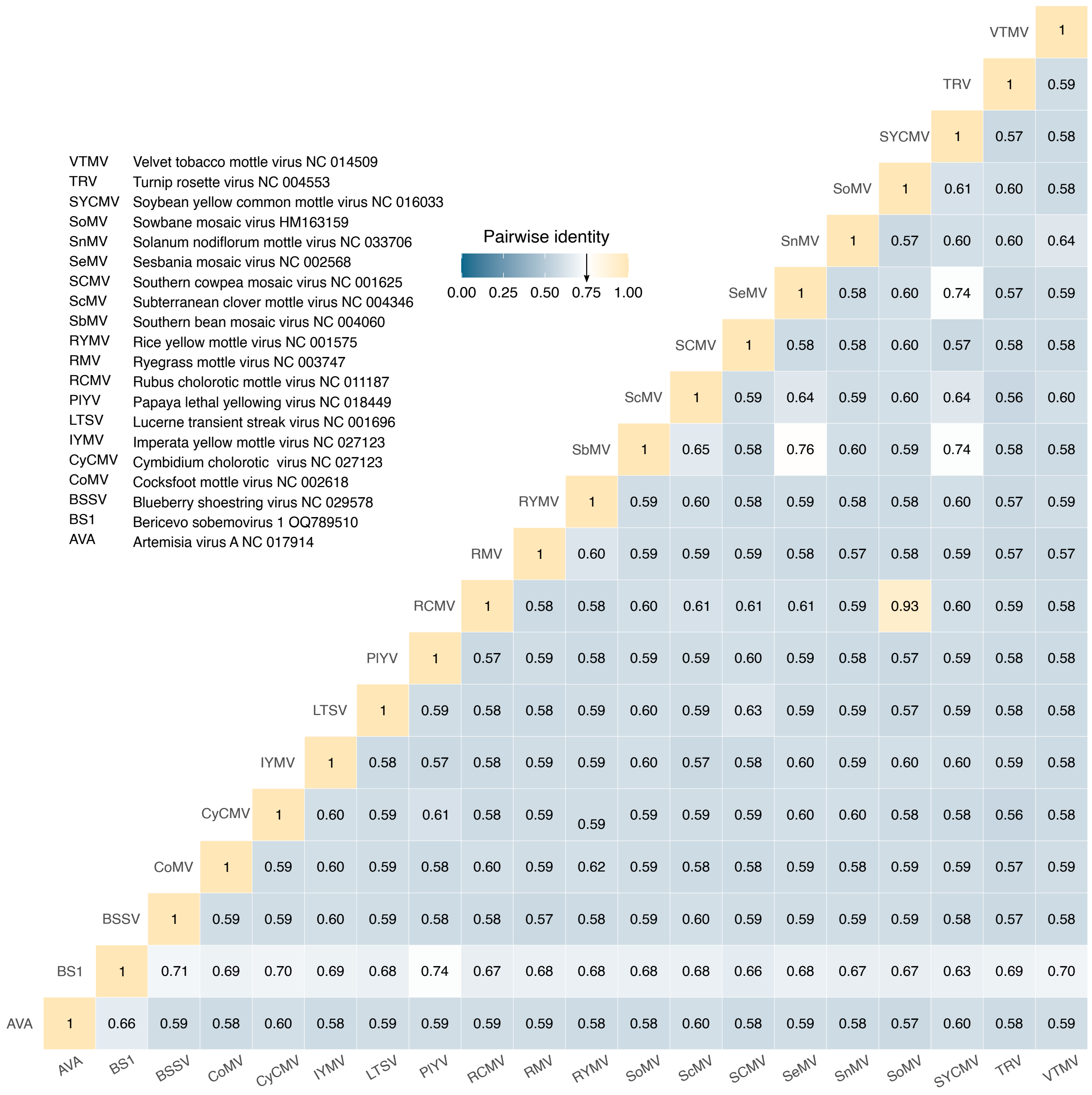

#### Supplementary Figure 7 - Pairwise similariy comparison of virus species from *Tobamovirus*

| | |
|---|---|
| ZGMMV | Zuchinni green mottle mosiac virus NC 003878 |
| YMV | Youcai mosaic virus NC 004422 |
| YtMMV | Yellow tallflower mild mottle virus NC 022801 |
| WMV | Wasabi mottle virus NC 003355 |
| TVV | Turnip vein-clearing virus NC 001873 |
| TsaMV | Tropical soda apple mosaic virus NC 030229 |
| ToMMV | Tomato mottle mosaic virus NC 022230 |
| ToMV | Tomato mosaic virus NC 002692 |
| ToBRFV | Tomato brown rugose fruit virus NC 028478 |
| TMV | Tobacco mosaic virus NC 001367 |
| TMGMV | Tobacco mild green mosaic virus NC 001556 |
| SFBV | Streptocarpus flower break virus NC 008365 |
| RMV | Ribgrass mosaic virus NC 002792 |
| ReMV | Rehmannia mosaic virus NC 009042 |
| PMV | Plumeria mosaic virus NC 026816 |
| PTV1 | Plantago tobamovirus 1 OL472109 |
| PMMoV | Pepper mild mottle virus NC 003630 |
| PfMV | Passionfruit mosiac virus NC015552 |
| PaMMV | Paprika mild mottle virus NC 004106 |
| ORV | Odontoglossum ringspot virus NC 001728 |
| OPV | Obuda pepper virus NC 003852 |
| KGMMV | Kyuri green mottle mosaic virus NC 003610 |
| HCSV | Hoya cholorotic spot virus NC 034509 |
| HLSV | Hibiscus latent Singapore virus NC 034509 |
| HLFFV | Hibiscus latent Fort Pierce virus NC 025381 |
| GT1 | Gorica tobamovirus 1 OQ789505 |
| FMV | Frangipani mosaic virus NC 014546 |
| CMV | Cucuber mottle virus NC 008614 |
| CGMMV | Cucumber green mottle mosaic virus NC 001801 |
| CFMMV | Cucumber fruit mottle mosaic virus NC 002633 |
| CYMV | Clitoria yellow mottle virus NC 016519 |
| CMMV | Cactus mild mottle virus NC 011803 |
| BMMV | Brugmansia mild mottle virus NC 010944 |
| BT1 | Bertoki tobamovirus 1 OQ789506 |
| BpMT | Bell pepper mottle tobamovirus NC 009642 |

Pairwise distance

0.00 0.25 0.50 0.75 1.00

0.90

| | BpMT | BT1 | BMMV | CMMV | CYMV | CFMMV | CGMMV | CMV | FMV | GT1 | HLFFV | HLSV | HCSV | KGMMV | OPV | ORV | PaMMV | PfMV | PMMOV | PTV1 | PMV | ReMV | RMV | SFBV | TMGMV | TMV | TBRFV | ToMV | ToMMV | TsaMV | TVV | WMV | YtfMMV | YMV | ZGMMV |
|---|---|---|---|---|---|---|---|---|---|---|---|---|---|---|---|---|---|---|---|---|---|---|---|---|---|---|---|---|---|---|---|---|---|---|---|
| ZGMMV | | | | | | | | | | | | | | | | | | | | | | | | | | | | | | | | | | | 1 |
| YMV | | | | | | | | | | | | | | | | | | | | | | | | | | | | | | | | | | 1 | 0.58 |
| YtMMV | | | | | | | | | | | | | | | | | | | | | | | | | | | | | | | | | 1 | 0.63 | 0.59 |
| WMV | | | | | | | | | | | | | | | | | | | | | | | | | | | | | | | | 1 | 0.64 | 0.84 | 0.58 |
| TVV | | | | | | | | | | | | | | | | | | | | | | | | | | | | | | | 1 | 0.83 | 0.63 | 0.82 | 0.58 |
| TsaMV | | | | | | | | | | | | | | | | | | | | | | | | | | | | | | 1 | 0.63 | 0.63 | 0.66 | 0.63 | 0.59 |
| ToMMV | | | | | | | | | | | | | | | | | | | | | | | | | | | | | 1 | 0.70 | 0.63 | 0.64 | 0.67 | 0.64 | 0.58 |
| ToMV | | | | | | | | | | | | | | | | | | | | | | | | | | | | 1 | 0.84 | 0.71 | 0.64 | 0.64 | 0.66 | 0.63 | 0.58 |
| ToBRFV | | | | | | | | | | | | | | | | | | | | | | | | | | | 1 | 0.81 | 0.81 | 0.71 | 0.63 | 0.63 | 0.66 | 0.63 | 0.59 |
| TMV | | | | | | | | | | | | | | | | | | | | | | | | | | 1 | 0.82 | 0.80 | 0.79 | 0.70 | 0.63 | 0.64 | 0.66 | 0.64 | 0.58 |
| TMGMV | | | | | | | | | | | | | | | | | | | | | | | | | 1 | 0.65 | 0.66 | 0.67 | 0.66 | 0.67 | 0.64 | 0.64 | 0.67 | 0.63 | 0.59 |
| SFBV | | | | | | | | | | | | | | | | | | | | | | | | 1 | 0.63 | 0.63 | 0.64 | 0.63 | 0.63 | 0.63 | 0.65 | 0.64 | 0.62 | 0.65 | 0.58 |
| RMV | | | | | | | | | | | | | | | | | | | | | | | 1 | 0.64 | 0.63 | 0.63 | 0.63 | 0.63 | 0.64 | 0.64 | 0.86 | 0.82 | 0.63 | 0.81 | 0.58 |
| ReMV | | | | | | | | | | | | | | | | | | | | | | 1 | 0.63 | 0.63 | 0.65 | 0.84 | 0.80 | 0.78 | 0.78 | 0.70 | 0.63 | 0.63 | 0.66 | 0.63 | 0.57 |
| PMV | | | | | | | | | | | | | | | | | | | | | 1 | 0.59 | 0.59 | 0.60 | 0.59 | 0.59 | 0.59 | 0.59 | 0.60 | 0.59 | 0.59 | 0.59 | 0.60 | 0.59 | 0.59 |
| PTV1 | | | | | | | | | | | | | | | | | | | | 1 | 0.59 | 0.63 | 0.86 | 0.64 | 0.63 | 0.63 | 0.63 | 0.64 | 0.64 | 0.63 | 0.85 | 0.81 | 0.63 | 0.80 | 0.58 |
| PMMoV | | | | | | | | | | | | | | | | | | | 1 | 0.64 | 0.59 | 0.70 | 0.63 | 0.63 | 0.67 | 0.70 | 0.70 | 0.71 | 0.70 | 0.79 | 0.63 | 0.64 | 0.66 | 0.63 | 0.58 |
| PfMV | | | | | | | | | | | | | | | | | | 1 | 0.60 | 0.60 | 0.59 | 0.60 | 0.60 | 0.59 | 0.59 | 0.59 | 0.58 | 0.60 | 0.58 | 0.61 | 0.59 | 0.60 | 0.60 | 0.59 | 0.59 |
| PaMMV | | | | | | | | | | | | | | | | | 1 | 0.60 | 0.67 | 0.63 | 0.59 | 0.66 | 0.63 | 0.62 | 0.66 | 0.67 | 0.67 | 0.67 | 0.67 | 0.67 | 0.63 | 0.64 | 0.70 | 0.64 | 0.58 |
| ORV | | | | | | | | | | | | | | | | 1 | 0.63 | 0.59 | 0.65 | 0.64 | 0.60 | 0.64 | 0.64 | 0.65 | 0.65 | 0.64 | 0.64 | 0.65 | 0.64 | 0.66 | 0.65 | 0.64 | 0.63 | 0.64 | 0.58 |
| OPV | | | | | | | | | | | | | | | 1 | 0.64 | 0.79 | 0.59 | 0.67 | 0.62 | 0.59 | 0.66 | 0.63 | 0.62 | 0.66 | 0.67 | 0.66 | 0.67 | 0.67 | 0.67 | 0.63 | 0.63 | 0.70 | 0.63 | 0.59 |
| KGMMV | | | | | | | | | | | | | | 1 | 0.58 | 0.58 | 0.59 | 0.59 | 0.59 | 0.58 | 0.58 | 0.59 | 0.59 | 0.58 | 0.59 | 0.57 | 0.59 | 0.59 | 0.58 | 0.59 | 0.58 | 0.59 | 0.58 | 0.59 | 0.83 |
| HCSV | | | | | | | | | | | | | 1 | 0.58 | 0.62 | 0.63 | 0.61 | 0.59 | 0.63 | 0.62 | 0.59 | 0.61 | 0.62 | 0.65 | 0.63 | 0.62 | 0.63 | 0.63 | 0.63 | 0.62 | 0.62 | 0.62 | 0.62 | 0.63 | 0.58 |
| HLSV | | | | | | | | | | | | 1 | 0.59 | 0.59 | 0.59 | 0.58 | 0.58 | 0.59 | 0.59 | 0.58 | 0.58 | 0.58 | 0.59 | 0.59 | 0.59 | 0.58 | 0.58 | 0.59 | 0.59 | 0.58 | 0.58 | 0.59 | 0.59 | 0.59 | 0.58 |
| HLFFV | | | | | | | | | | | 1 | 0.71 | 0.58 | 0.60 | 0.59 | 0.59 | 0.59 | 0.60 | 0.59 | 0.59 | 0.59 | 0.58 | 0.59 | 0.58 | 0.60 | 0.59 | 0.58 | 0.58 | 0.58 | 0.59 | 0.59 | 0.59 | 0.58 | 0.59 | 0.59 |
| GT1 | | | | | | | | | | 1 | 0.58 | 0.58 | 0.63 | 0.58 | 0.62 | 0.65 | 0.62 | 0.59 | 0.63 | 0.69 | 0.59 | 0.62 | 0.69 | 0.67 | 0.63 | 0.62 | 0.63 | 0.64 | 0.63 | 0.64 | 0.69 | 0.69 | 0.63 | 0.68 | 0.58 |
| FMV | | | | | | | | | 1 | 0.59 | 0.59 | 0.59 | 0.59 | 0.59 | 0.60 | 0.59 | 0.59 | 0.59 | 0.60 | 0.59 | 0.73 | 0.59 | 0.59 | 0.59 | 0.59 | 0.60 | 0.59 | 0.59 | 0.59 | 0.60 | 0.59 | 0.59 | 0.59 | 0.59 | 0.59 |
| CMV | | | | | | | | 1 | 0.59 | 0.58 | 0.59 | 0.59 | 0.58 | 0.64 | 0.60 | 0.59 | 0.59 | 0.59 | 0.58 | 0.59 | 0.58 | 0.59 | 0.58 | 0.58 | 0.59 | 0.59 | 0.59 | 0.59 | 0.59 | 0.58 | 0.59 | 0.58 | 0.59 | 0.59 | 0.65 |
| CGMMV | | | | | | | 1 | 0.65 | 0.58 | 0.58 | 0.58 | 0.59 | 0.58 | 0.65 | 0.59 | 0.59 | 0.59 | 0.60 | 0.59 | 0.59 | 0.58 | 0.59 | 0.59 | 0.59 | 0.58 | 0.58 | 0.59 | 0.58 | 0.58 | 0.59 | 0.60 | 0.59 | 0.58 | 0.60 | 0.64 |
| CFMMV | | | | | | 1 | 0.65 | 0.64 | 0.58 | 0.58 | 0.61 | 0.58 | 0.59 | 0.74 | 0.59 | 0.60 | 0.59 | 0.59 | 0.59 | 0.58 | 0.59 | 0.57 | 0.58 | 0.59 | 0.58 | 0.59 | 0.58 | 0.58 | 0.58 | 0.58 | 0.59 | 0.59 | 0.60 | 0.58 | 0.73 |
| CYMV | | | | | 1 | 0.58 | 0.58 | 0.59 | 0.58 | 0.59 | 0.57 | 0.57 | 0.58 | 0.58 | 0.58 | 0.59 | 0.58 | 0.60 | 0.58 | 0.58 | 0.58 | 0.58 | 0.58 | 0.58 | 0.58 | 0.58 | 0.58 | 0.59 | 0.59 | 0.57 | 0.59 | 0.59 | 0.58 | 0.57 | 0.58 |
| CMMV | | | | 1 | 0.56 | 0.57 | 0.58 | 0.57 | 0.59 | 0.58 | 0.58 | 0.58 | 0.58 | 0.57 | 0.58 | 0.57 | 0.58 | 0.59 | 0.58 | 0.57 | 0.57 | 0.58 | 0.58 | 0.59 | 0.60 | 0.58 | 0.58 | 0.58 | 0.58 | 0.58 | 0.59 | 0.58 | 0.58 | 0.57 | 0.58 |
| BMMV | | | 1 | 0.58 | 0.58 | 0.59 | 0.59 | 0.59 | 0.58 | 0.63 | 0.58 | 0.59 | 0.62 | 0.59 | 0.67 | 0.64 | 0.68 | 0.60 | 0.67 | 0.63 | 0.58 | 0.67 | 0.63 | 0.63 | 0.67 | 0.67 | 0.67 | 0.67 | 0.67 | 0.67 | 0.63 | 0.63 | 0.67 | 0.63 | 0.59 |
| BT1 | | 1 | 0.63 | 0.58 | 0.58 | 0.58 | 0.58 | 0.58 | 0.59 | 0.69 | 0.59 | 0.59 | 0.62 | 0.58 | 0.63 | 0.64 | 0.63 | 0.60 | 0.63 | 0.87 | 0.59 | 0.62 | 0.86 | 0.64 | 0.64 | 0.63 | 0.62 | 0.63 | 0.63 | 0.63 | 0.87 | 0.82 | 0.62 | 0.81 | 0.59 |
| BpMT | 1 | 0.63 | 0.68 | 0.59 | 0.58 | 0.59 | 0.59 | 0.58 | 0.59 | 0.64 | 0.59 | 0.59 | 0.63 | 0.59 | 0.67 | 0.65 | 0.67 | 0.59 | 0.70 | 0.64 | 0.59 | 0.74 | 0.65 | 0.63 | 0.67 | 0.76 | 0.76 | 0.78 | 0.77 | 0.71 | 0.64 | 0.64 | 0.66 | 0.63 | 0.59 |

Supplementary Figure 8 - Pairwise similariy comparison of detected plantago tobamovirus 1 on two selected regious around coat protein (left) and RdRp (right)

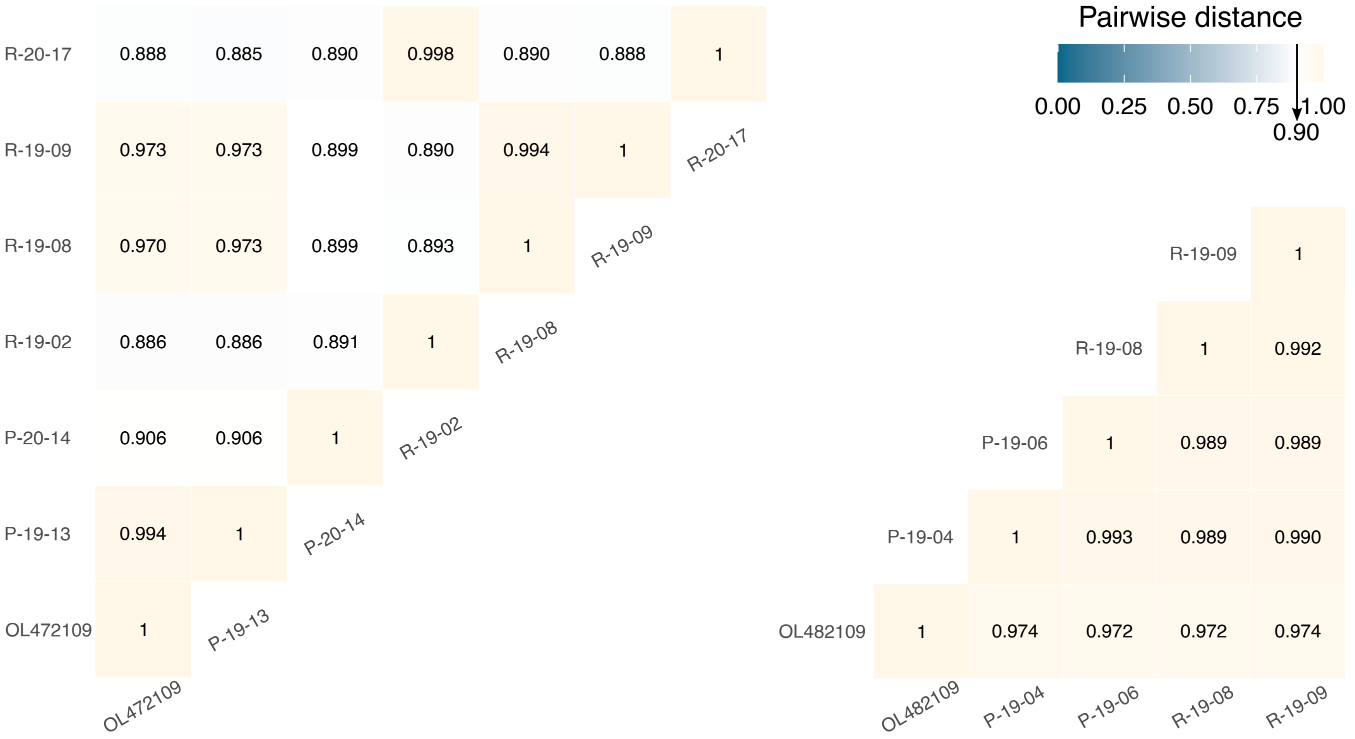


Supplementary Figure 9 - Results of RT-qPCR detection for pepper mild mottle virus and tomato mosaic virus for two sampling years (reads divided due to subsampling, as indicated in Supplementary Information 1). Smaller dot represents the resuts before concentration and larger dot after concentration of the same sample. Lenght of the line indicates the effect of concentration expressed as decrease in obtained Cq.

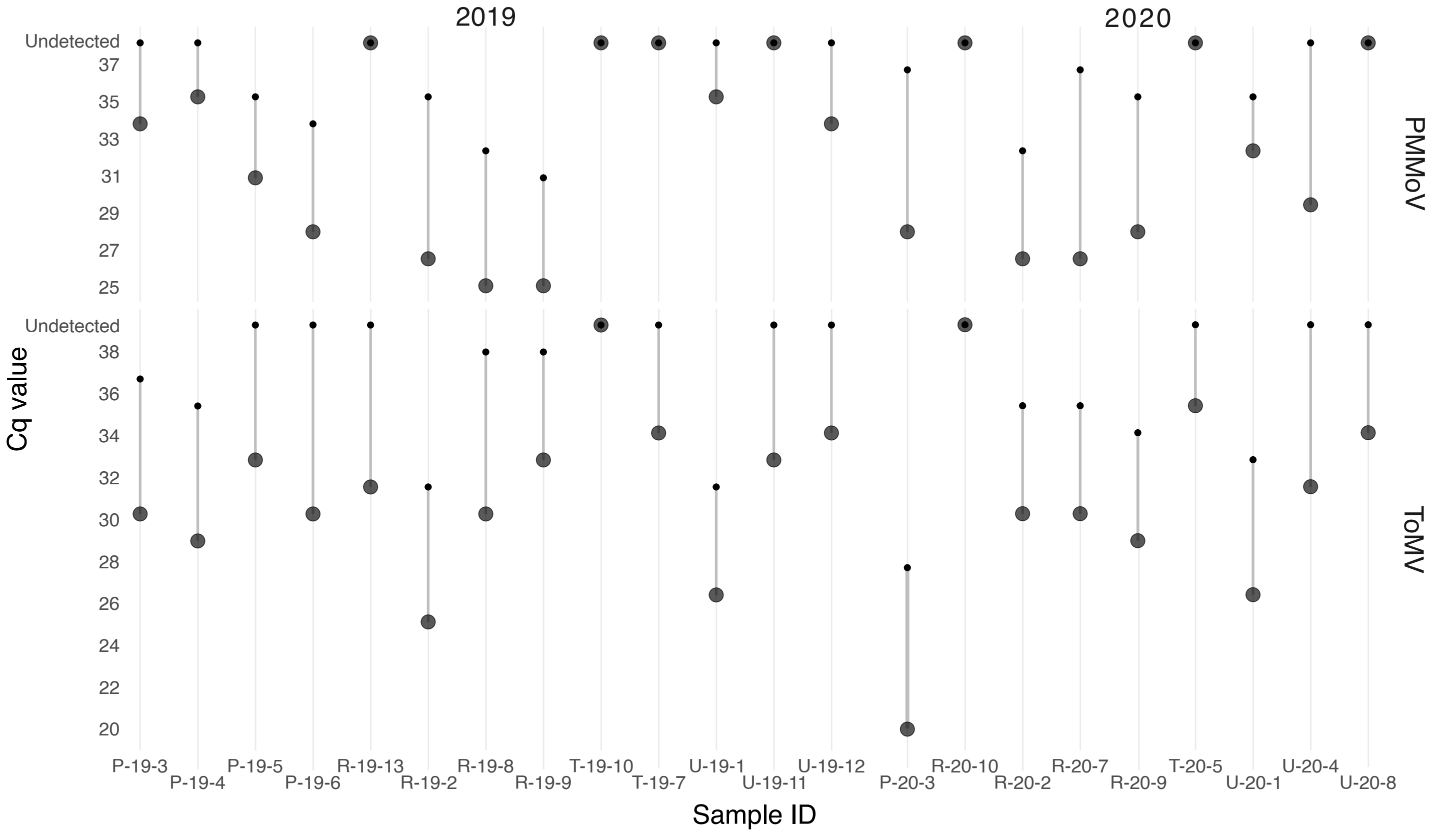

Supplementary figure 10-12 - Phylogenetic trees show relationship of known and novel (in bold) virus species for a given genus. GenBank accession numbers are are indicated next to the sequence for the known viruses or can be found in Supplementary information 1 for nover sequences. Details regarding the phylogenetic analysis are in Supplementary information 1.

Supplementary figure 10 - maximum likelihood phylogenetic tree based on the alingment of conserved RdRp protein sequence of virus species, including new sequence, under *Aureusvirus*

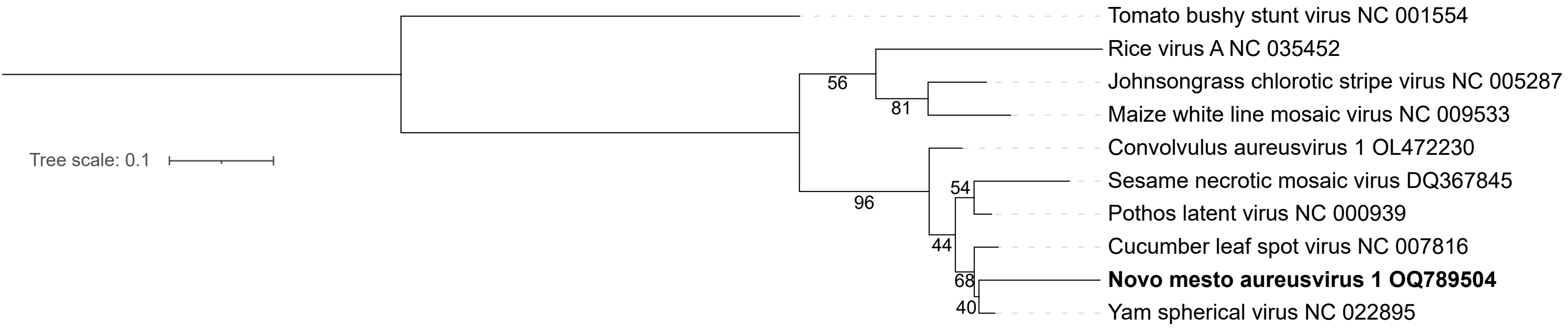


Supplementary figure 11 - maximum likelihood phylogenetic tree based on the alingment of conserved RdRp protein sequence of virus species, including new sequence, under *Betanecrovirus*

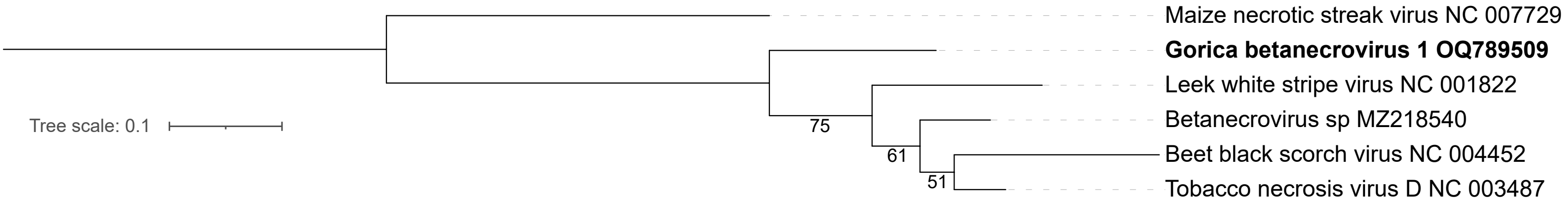


Supplementary figure 12 - maximum likelihood phylogenetic tree based on the alingment of whole nucleotide sequence of virus species, including new sequence, under *Sobemovius*

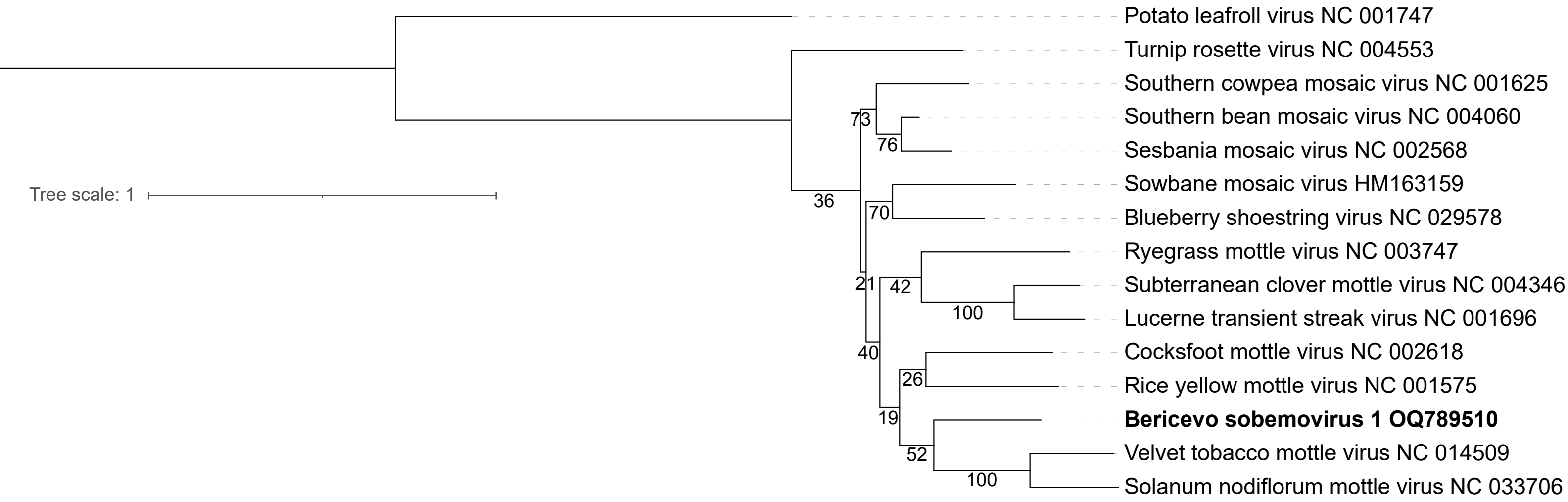
